# A combination of convergent extension and differential adhesion explains the shapes of elongating gastruloids

**DOI:** 10.1101/2023.05.24.541949

**Authors:** Martijn A. de Jong, Esmée Adegeest, Noémie M. L. P. Bérenger-Currias, Maria Mircea, Roeland M. H. Merks, Stefan Semrau

## Abstract

Gastruloids have emerged as highly useful *in vitro* models of mammalian gastrulation. One of the most striking features of 3D gastruloids is their elongation, which mimics the extension of the embryonic anterior-posterior axis. Although axis extension is crucial for development, the underlying mechanism has not been fully elucidated in mammalian species. Gastruloids provide an opportunity to study this morphogenic process *in vitro*. Here, we measure and quantify the shapes of elongating gastruloids and show, by Cellular Potts model simulations, that a combination of convergent extension and differential adhesion can explain the observed shapes. We reveal that differential adhesion alone is insufficient and also directly observe hallmarks of convergent extension by time-lapse imaging of gastruloids. Finally, we show that gastruloid elongation can be abrogated by inhibition of the Rho kinase pathway, which is involved in convergent extension *in vivo*. All in all, our study demonstrates, how gastruloids can be used to elucidate morphogenic processes in embryonic development.

## Introduction

Embryonic morphogenesis produces a bewildering diversity of shapes using a range of intricate mechanisms. Over many decades, through careful observation and perturbation of live embryos, some of these mechanisms have been elucidated, while many morphogenic processes remain ill-understood. Mammalian embryogenesis is particularly challenging to study, because crucial morphogenic events occur *in utero*. For example, gastrulation, the process that transforms the mammalian embryo from a single epithelial layer (the epiblast) into three distinct germ layers (ectoderm, endoderm and mesoderm), follows implantation of the embryo in the uterine wall. Gastrulation is of crucial importance, because the germ layers are precursors of specific tissues and hence indispensable for proper organogenesis. A highly coordinated sequence of differentiation and cell movements accompanies not only germ layer formation but also the subsequent establishment of the body axes and embryo elongation. Recent advances in light sheet microscopy (1; 2) and *ex vivo* culture of embryos (3; 2) have started to shed more light on these processes but the sheer complexity of the system and limited opportunities to manipulate or perturb it, make it very difficult to pinpoint specific mechanisms.

Recently, stem cell-derived *in vitro* systems have emerged as accessible models that aim to mimic developmental processes with a minimal number of necessary components. These models have the advantage of being easy to produce at scale and are amenable to perturbation. Their reduced complexity, compared to an actual embryo, allows us to study specific developmental processes in isolation. Gastruloids, established by van den Brink et al. (4), are aggregates of mouse embryonic stem cells (mESCs) that exhibit hallmarks of gastrulation. Strikingly, gastruloids elongate during the last 24 h of a 4 day differentiation trajectory. This elongation is thought to mimic the formation and subsequent extension of the first body axis in the mouse embryo: the anterior-posterior (AP) axis. This axis is formed at the onset of gastrulation in the posterior pole of the embryo. In a specific region, the primitive streak, cells extensively ingress and migrate, which eventually leads to the formation of the AP axis. While AP axis elongation has been studied extensively in non-mammalian vertebrates (5), much less is known about the mechanical cell interactions and related movement patterns that shape the mouse or human embryo during AP axis formation. Elongating gastruloids thus present an exciting opportunity to reveal those mechanisms (6).

Tissue elongation can be driven by a range of different mechanisms, including directed cell migration as well as localized proliferation or cell growth (5). Early elongation of the AP axis in Drosophila (7), Xenopus (8) and zebrafish (9), involves yet another mechanism: convergent extension (CE). Cells of the AP axis initially stretch in one direction and then intercalate, such that the tissue elongates in the perpendicular direction. Importantly, CE does not require cell division. Two related mechanisms have been observed to underlie CE. The “crawling” mechanism involves mediolaterally-oriented protrusions on the poles of the cells. These protrusions generate integrin-dependent forces on the extracellular matrix and adjacent cells. By contrast, the “junction contraction” mechanism employs active contraction of cellular interfaces to drive intercalation. Evidence in *Drosophila* (10), *Xenopus* (11) and mouse (12) suggests that the two mechanisms may act in concert, but are regulated independently (13). In *Drosophila*, in absence of either of the two mechanisms (10), CE is slowed down but not stopped completely. In a recent combined experimental and mathematical modeling study in *Xenopus* (11) concurrent crawling and contraction was demonstrated by live imaging. A mathematical model of CE revealed that both mechanisms occurring together results in stronger intercalation compared to either of the two occurring independently. Thus, the “crawling” and “junction contraction” mechanisms may reinforce each other but suffice to drive CE independently. While live imaging has established a role for CE in mouse embryogenesis, specifically in neural tube formation (14; 12), its involvement in AP axis elongation is much less established.

In this manuscript we will demonstrate an alternative way to test whether CE underlies elongation. We reasoned that the shape of elongating gastruloids should reflect the mechanism that causes their elongation. To test this idea we imaged gastruloids at multiple time points and quantified their shapes using Lobe-Contribution Elliptic Fourier Analysis (LOCO-EFA) (15). To understand how the observed shapes might be created by different types of cell interactions we adopted an *in silico* approach based on the Cellular Potts model (CPM) (16; 17) aka Glazier-Graner-Hogeweg model. The CPM represents single cells as sets of connected sites on a lattice. Cells move and change their shape by exchanging lattice sites with neighboring cells. These changes in site occupancy are generated randomly using a Monte Carlo method. The probability of a change is governed by the total energy of the system, which is minimized in the course of the simulation. Specific terms in the expression for the total energy (the Hamiltonian) can be used to model interactions between cells, such as non-uniform adhesion along cell boundaries (18) or active pulling mediated by filopodia (19). We first used this *in silico* model to show that cell-to-cell variability in adhesion is not sufficient to explain the observed shapes. We then adopted an unbiased, filopodial tension model of CE (19) based on the “crawling” mechanism, and used the experimental measurements to constrain the free model parameters. The simulations showed that CE alone can reproduce much of the observed shape variety. By introducing multiple cell types in our simulation, we could further improve the agreement with the experiments. To confirm the role of CE experimentally we pursued live-imaging of elongating gastruloids and observed hallmarks of CE. Finally, we demonstrated that inhibition of the Rho kinase (ROCK) pathway, which was found to be necessary for CE *in vivo*, impairs gastruloid elongation.

## Materials and methods

### Experimental Methods

#### Tissue culture

##### Cell lines

Gastruloids were generated from E14 mouse ES cells. The E14 cell line was provided by Alexander van Oudenaarden. To visualize cell membranes in gastruloids, a cell line was created that expresses the fluorescent reporter protein mCherry fused to glycosylphosphatidylinositol (GPI). The fused GPI anchors the mCherry protein to the cell membrane. The mCherry-GPI cell line was created by introducing a GPI:mCherry transgene in the E14 cell line. Both cell lines were routinely cultured in KnockOut DMEM medium (Gibco) supplemented with 10% ES certified fetal bovine serum (US origin, Gibco), 0.1 mM 2-Mercaptoethanol (Sigma-Aldrich), 1 x 100 U/mL penicillin/streptomycin, 1 x MEM Non-Essential Amino Acids (Gibco), 2 mM L-glutamine (Gibco) and 1000 U/mL recombinant mouse LIF (ESGRO). Cells were grown in tissue culture-treated dishes that were pre-coated with 0.2% gelatin for a minimum of 10 min at 37°C. Cells were passaged three times a week at a 1:6 ratio for up to 20 passages. Cell cultures were maintained at 37°C and 5% CO_2_.

##### Gastruloid culture

To grow gastruloids, we followed the protocol originally described by van den Brink et al. (4). ES cells were collected from the dish after dissociation with Trypsin-EDTA (0.25%) for 3 min at 37°C. Trypsinization was stopped by adding an equal amount of culture medium, followed by gentle trituration with a pipet and centrifugation (300 rcf, 3 min). The cell pellet was resuspended in 2 mL of freshly prepared N2B27 medium: DMEM/F12 with L-glutamine and sodium bicarbonate, without HEPES (Sigma, D8062) supplemented with 0.5 x N2 supplement (Gibco), 0.5 x B27 supplement (Gibco), 0.5 mM L-glutamine (Gibco), 0.5 x MEM Non-Essential Amino Acids (Gibco), 0.1 mM 2-Mercaptoethanol (Sigma-Aldrich) and 1 x 100 U/mL penicillin/streptomycin. Cells were counted to determine the cell concentration. A total of 200 cells were seeded in 40 *µ*L of N2B27 medium per well in a U-bottom low-adherence 96-well plate (Greiner CELLSTAR). After 48 h, the resulting aggregates were exposed to a 24 h pulse of GSK-3 inhibitor by adding 150 *µ*L N2B27 medium supplemented with 3 *µ*M CHIR (CHIR99021, Axon Medchem) to each well. At 72 h after cell seeding, the 24 h CHIR pulse was stopped by replacing 150 *µ*L of the medium in each well with fresh N2B27 medium. At 96 h after seeding, gastruloids were collected and fixed to analyze their shapes. For Fig. 1, gastruloids were also collected at 48 h and 72 h after cell seeding. For the time lapses, we grew mosaic gastruloids by seeding a mixture of E14-GPI:mCherry ES cells and regular E14 ES cells at a ratio of 1:16. For the experiments with the ROCK inhibitor, 10 *µ*M of Y-27632 (Sigma) was added to the N2B27 medium at 72 h after cell seeding. Treatment with Y-27632 was continued for 24 h, up until 96 h after cell seeding. The control group received N2B27 medium without Y-27632 at 72 h after cell seeding. We performed a total of two biological replicates for this experiment.

**Fig 1.**
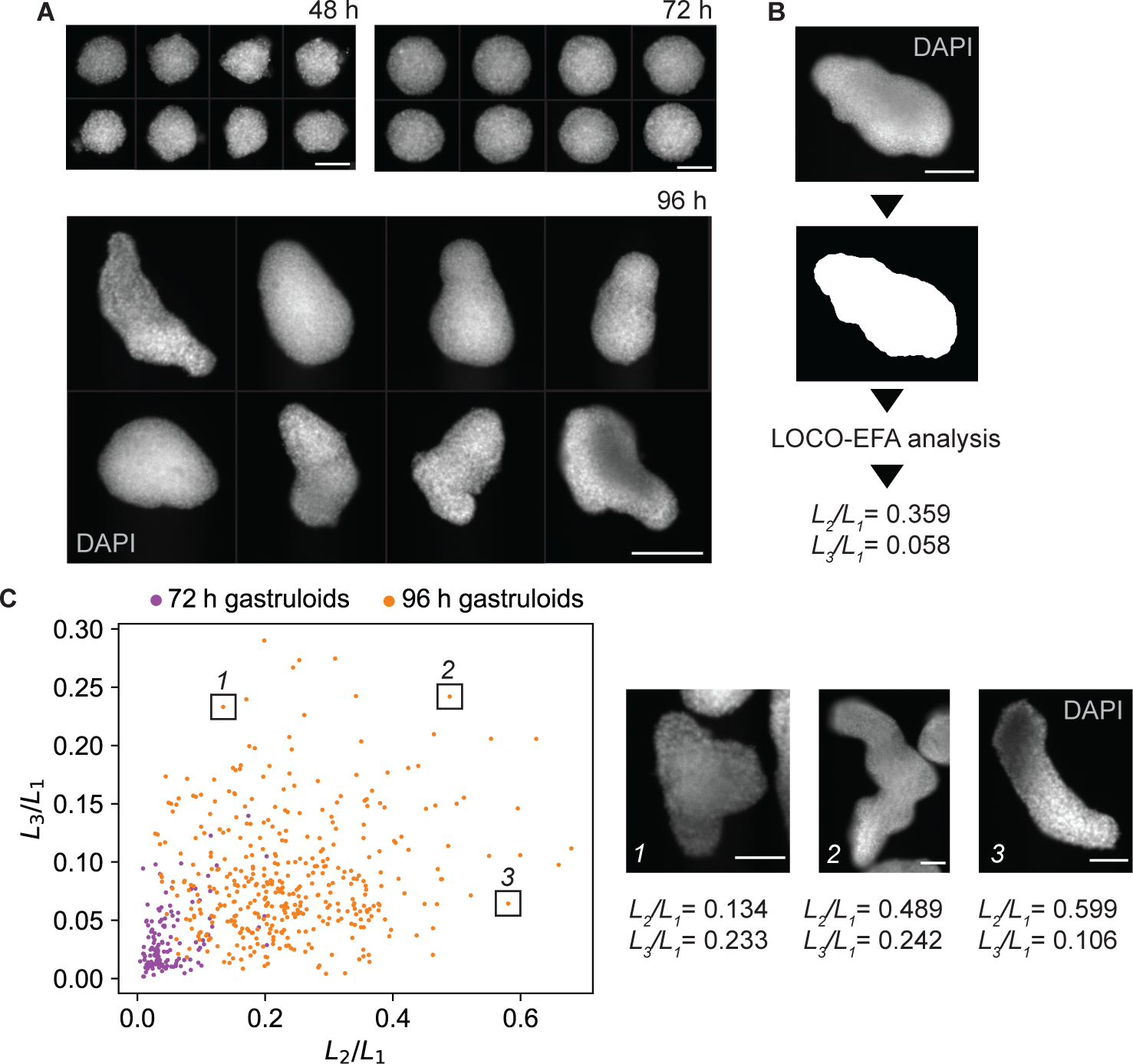
Experimental observation of gastruloids and shape quantification using LOCO-EFA. A: Representative images of fixed gastruloids after 48 h, 72 h and 96 h. Cell nuclei were stained with DAPI. Shown is the mid-plane of a z-stack. Scale bar: 100 *µ*m (48 h and 72 h), 200 *µ*m (96 h). B: Image of a gastruloid with binary mask created from that image. Numerical values of *L*_2_*/L*_1_ and *L*_3_*/L*_1_ for this gastruloid are given. Scale bar: 100 *µ*m. C: LOCO-EFA analysis of measured gastruloids at 72 h (n = 132, two biological replicates) and 96 h (n = 413, 8 biological replicates). Each data point is a gastruloid. The scatter plot shows the scaled LOCO-EFA coefficients *L*_2_*/L*_1_ and *L*_3_*/L*_1_. The 3 insets show images of the gastruloids highlighted in the LOCO-EFA scatter plot. Cell nuclei were stained with DAPI. Shown is the mid-plane of a z-stack. For each gastruloid, the scaled LOCO-EFA coefficients *L*_2_*/L*_1_ and *L*_3_*/L*_1_ are given. Scale bar: 100 *µ*m.

#### Immunostaining

##### Fixation and blocking

Gastruloids were fixed by incubating them in 4% paraformaldehyde (PFA, Alfa Aeser) at 4°C for 4 h. Fixation of gastruloids was stopped by washing them three times in PBS supplemented with 1% bovine serum albumin (BSA). Gastruloids were incubated in blocking buffer (PBS, 1%BSA, 0.3% Triton-X-100) at 4°C for a minimum of 1 h prior to immunostaining.

##### Whole-mount immunolabeling

Immunolabeling of gastruloids was based on the protocol described by Dekkers et al. (20). In short, after fixation and blocking, gastruloids were incubated with primary antibodies in organoid washing buffer (OWB) (PBS, 0.2% BSA, 0.1% Triton-X-100) supplemented with 0.02% sodium dodecyl sulfate (SDS), referred to as OWB-SDS. Incubation was carried out at 4°C overnight on a rolling mixer (30 rpm). The following primary antibodies were used: rat anti-SOX2 (1:200, 14-9811-82, Thermo Fisher Scientific), goat anti-T (1:200, AF2085, R&D Systems) and rat anti-Cerberus1 (1:200, MAB1986, R&D Systems). The next day, gastruloids were washed in OWB-SDS on a tube rotator for 2 h at room temperature (RT). This step was repeated for a total of three times. After washing, gastruloids were incubated with secondary antibodies and 4’,6-diamidino-2-phenylindole (DAPI, 1 *µ*g/mL, Merck) in OWB-SDS at 4°C overnight, in the dark, on a rolling mixer (30 rpm). The following secondary antibodies were used: donkey anti-rat Alexa Fluor 488 (1:400, A-21208, Thermo Fisher Scientific), donkey anti-goat Alexa Fluor 555 (1:400, A-21432, Thermo Fisher Scientific), donkey anti-goat Alexa Fluor 488 (1:400, A-11055, Thermo Fisher Scientific) and chicken anti-rat Alexa Fluor 647 (1:400, A-21472, Thermo Fisher Scientific). The next day, gastruloids were washed again three times with OWB-SDS on a tube rotator for 2 h at RT. After washing, gastruloids were immediately imaged or stored at 4°C prior to imaging.

##### Cryo-sectioning and immunolabeling of sections

To prepare the gastruloids for cryo-sectioning, gastruloids were first sequentially incubated in sucrose solutions (10%, 20% and 30% in PBS) for 30 min at 27°C. After these sucrose incubation steps, the gastruloids were transferred to a Tissue-TEK Cryomold 10*×* 10 *×* 5 mm (Sakura). There, the 30% sucrose solution was removed and replaced with optimal cutting temperature (O.C.T.) compound (VWR). The O.C.T.-filled cryomolds were rapidly frozen on dry ice and stored at −80°C until sectioning. On the day of sectioning, the cryomolds were transferred to the cryostat to let them reach the cutting temperature of −20°C. After about one hour, the cryomolds were removed and the O.C.T. blocks were mounted on the cryostat. Sections with a thickness of 10 *µ*m were cut and placed on Epredia^TM^ SuperFrost Plus^TM^ Adhesion slides (Merck). Slides were stored at −80°C until staining. To perform immunofluorescence staining on the sections, slides were thawed at RT and subsequently washed with PBS for 10 min to dissolve the O.C.T.. After washing, the sections were incubated overnight at 4°C with the following primary antibodies diluted in blocking buffer: rat anti-SOX2 (1:200, 14-9811-82, Thermo Fisher Scientific), goat anti-T (1:200, AF2085, R&D Systems) and rabbit anti-KI-67 (1:200, MA5-14520, Thermo Fisher Scientific). The next day, slides were washed twice for 10 min in PBS at RT. After washing, slides were incubated for 4 h at 4°C with DAPI (1 *µ*g/mL, Merck) and the following secondary antibodies diluted in blocking buffer: donkey anti-rat Alexa Fluor 488 (1:400, A-21208, Thermo Fisher Scientific), donkey anti-goat Alexa Fluor 555 (1:400, A-21432, Thermo Fisher Scientific) and donkey anti-rabbit Alexa Fluor 647 (1:200, A-31573, Thermo Fisher Scientific). After 4 h, slides were washed three times with PBS for 10 min at RT. Finally, sections were mounted in ProLong Gold^TM^ Antifade Mountant (Thermo Fisher Scientific). Slides were cured for 48 h at RT prior to imaging.

#### Imaging

##### Imaging of fixed gastruloids

For imaging wholemount immunostained gastruloids, gastruloids were transferred to *µ*-slide 8 well glass bottom chambers (ibidi, Cat.No 80827). Both wholemount gastruloids and cryo-sectioned gastruloids were imaged on a Nikon Ti-Eclipse epifluorescence microscope equipped with an Andor iXON Ultra 888 EMCCD camera and dedicated, custom-made fluorescence filter sets (Nikon). Images were taken using primarily a 10*×*/0.3 Plan Fluor DLL objective (Nikon) and a 20*×*/0.5 Plan Fluor DLL objective (Nikon). z-stacks of whole-mount gastruloids were collected with a 10 *µ*m distance between planes. The images of the immunostained cryo-sections were pre-processed in ImageJ to remove unspecific background signal, which was present in the Brachyury/T and Sox2 channels. For these channels, the background was subtracted using the rolling ball function (radius: 100 pixels = 64 *µ*m).

##### Time-lapse imaging

A time lapse of the mosaic gastruloid during the final hours of elongation was performed between 91 and 96 h after cell seeding. Imaging was performed with a home-built light-sheet fluorescence microscope that kept the temperature and CO_2_ level constant during the experiment at 37°C and 5%, respectively. During the time lapse, z-stacks were taken with a step size of 4 *µ*m with a total of 40 planes per time point, thereby covering an imaging depth of 160 *µ*m. Each image was taken with a 160 ms exposure time, and a z-stack was taken every 10 min for a total of 5 hours. The mCherry fluorescent protein was excited by a 561 nm laser with a laser power of 50 *µ*W (measured at the image plane). With a light-sheet thickness of 8.7 *µ*m, determined by the thickness of the beam (full width half maximum,FWHM), the irradiance at the sample plane was calculated to be 79 W/cm^2^.

##### Design of the light-sheet microscope

To perform the time lapse imaging of the mosaic gastruloid, we used a home-built light-sheet fluorescence microscope. This microscope is based on the design described in (21) and some enhancements towards organoid imaging (22). In short, the setup consists of an illumination arm, an imaging arm and a custom specimen chamber. The illumination arm of the setup is equipped with a C-FLEX laser combiner (Hübner Photonics) that combines three diode lasers (405 nm, 488 nm, 647 nm; 06-01 series, Cobolt) and one diode-pumped solid state laser (561 nm; 06-01 series, Cobolt) into a single path through a single-mode/polarization-maintaining optical fiber (kineFLEX, Qioptic). The output of the fiber is coupled to an achromatic collimator (PAF2-A4A, Thorlabs) to collimate the outcoming beam. The collimated beam is first passed through a neutral density filter wheel, set to an attenuation factor of 10*×*, to keep the laser power at the image plane low. The attenuated beam then enters a two-axis deflection unit (MINISCAN-II-10, Raylase) whose two galvanometric scanning mirrors are controlled by a desktop device (ScanMaster SM1000, Cambridge Technology). A telecentric F-Theta lens (S4LFT0061/065, Sill Optics) is connected to the output of the deflection unit to ensure that angular deflections of the beam by the mirrors are translated into a linear displacement in the specimen. Scanning the beam along one axis therefore results in a virtually-scanned light-sheet. After the scan head, the beam is sent to two kinematic mirrors that make up a periscope to bring the beam to the height of the specimen chamber. The beam then travels through a tube lens (*f* = 200 mm, Nikon) to obtain a collimated beam. The tube lens is placed in front of the illumination objective such that its focal plane coincides with the illumination objective’s back focal plane. A 10*×* magnification water-dipping objective (CFI Plan Fluor 10X W, 0.3 NA, 16 mm W.D., Nikon) is orientated horizontally and projects the virtually-scanned light-sheet onto the sample plane. The light-sheet thickness was measured to be 8.7 *µm* (FWHM). In the imaging arm, the emitted fluorescence is collected from below by a 25*×* magnification water-dipping objective (CFI75 Apochromat 25XC W, 1.1 NA, 2 mm W.D., Nikon). A kinematic mirror positioned below the detection objective directs the emitted fluorescence through an emission filter, placed inside a motorized filter wheel (FW103H/M, Thorlabs). A 605/15 emission filter (FF01-605/15-25, Semrock) was used to collect the emitted fluorescence from the mCherry-GPI cells. The filtered light then travels through a tube lens (*f* = 200 mm, Nikon) that projects the image onto a sCMOS camera (Iris 15, Teledyne Photometrics). The illumination and detection objectives are mounted in a customized specimen chamber. In this chamber, the temperature is regulated by an external water bath (CORIO CD-B13, Julabo) that continuously supplies water at given temperature. A drain in the specimen chamber at the height of the sample holder returns the excess water to the water bath. The CO_2_ level inside the specimen chamber is controlled by a CO_2_ sensor and controller (ProCO2 P120, Biospherix). The sample holder is moved by three nanometer linear-positioners (SLC-24, SmarAct) along the x, y and z-axis. The specimen is placed inside a disposable, ready-to-use, sterile *TruLive3D* dish (TruLive3D, Bruker). Two of the dishes can be placed in a customized sample holder. The transparent FEP foil of the TruLive3D dish has a curved shape, but a refractive index matching that of water. It therefore separates the cell culture medium inside the dish from the water that is outside in the specimen chamber without introducing refraction.

### Computational Methods

#### Shape analysis

##### Shape extraction from images

To enable the quantification of gastruloid shapes, wide-field images of fixed, DAPI-stained gastruloids were collected and processed. To extract shapes, we developed a computational tool that segments shapes, often multiple in one image, into individual binary masks. The segmentation steps are described below. First, we applied a Gaussian filter to the images with a standard deviation of 5 pixels. Then, we found a global threshold with Otsu’s method (23). We applied the threshold to the filtered image and subsequently applied both dilation and opening with a radius of 10 pixels to the resulting binary image to avoid holes in the segmented objects. Next, we calculated average Euclidean distances from each foreground pixel to the closest background pixel and identified local maxima in the distance matrix. Local maxima with a minimal distance of 70 pixels were chosen as markers for each object in the image. The background was added as an additional marker. Furthermore, we used the Scharr transform (24) on the filtered image to generate an elevation map. Then, we applied the watershed algorithm (25) on the elevation map using the markers as starting points. Objects that had a size of less than 1000 pixels were excluded, as well as objects touching the border of the image.

##### Lobe-Contribution Elliptic Fourier Analysis

Due to the complexity of gastruloid shapes we found scalar shape metrics to be insufficient to adequately describe the observed shape distributions. Hence, we resorted to a recent improvement on classical Fourier Analysis methods to describe the shapes. Lobe-Contribution Elliptic Fourier Analysis (LOCO-EFA) (15) was developed for the quantification of cell shapes but is applicable to arbitrary two-dimensional, compact shapes. The LOCO-EFA coefficient *L_n_* quantifies the contributions of a mode with *n* lobes to the two-dimensional shape. *L*_1_ is thus essentially a measure for the linear size of the shape. To make the experimental data (which is measured in physical spatial units) comparable with the simulations (which are run on a lattice without specification of a physical size), all values of *L_n_* (for *n ≥* 2) were scaled by dividing by *L*_1_ of the respective shape. For simulated shapes, average LOCO-EFA coefficients were calculated from 100 independent simulations. For experimental shapes, LOCO-EFA coefficients were calculated from gastruloids originating from 2 biological replicates (72 h time point, n = 132) or 8 biological replicates (96 h time point, n = 413).

#### Analysis of mosaic gastruloids

##### Time lapse: image denoising

Before analysis, images of the time lapse of the mosaic mCherry-GPI gastruloid were cropped in Fiji (26) around the gastruloid to reduce the file size. Next, z-stacks of each time point were denoised using the Noise2Void (N2V) machine learning-based method (27). N2V trains directly on the data to be denoised and does not require noisy image pairs or clean target images. Our time lapse data set consisted of 31 z-stacks, with each z-stack containing 40 image planes with a (cropped) area of 2960 *×* 1878 pixels. In total, 10672 non-overlapping 3D patches of size 32*×* 64 *×* 64 pixels (z, y, x) were extracted from a single z-stack. 90% of the patches were used for training and 10% were used for prediction. Training a model on one of these z-stacks on one GPU, using 200 epochs and 75 trains steps per epoch, took roughly 4 hours. A trained model can be used to denoise data that was not used for training, but was recorded with the same image settings. Here it is important that the data set has a similar noise distribution. Throughout our time lapse, we detected fluctuations in the noise distribution. We therefore trained three different models: the first model was trained on the 1*^st^* time point and was used to predict denoised images from the z-stack of this time point only; the second model was trained on the 7*^th^* time point and was used to predict denoised images from z-stacks of the 2*^nd^* to 7*^th^* time point; the third model was trained on the last time point and was used to predict denoised images from z-stacks of the 8*^th^* to the last (31*^st^*) time point. Examples of raw and denoised images for the three models are displayed in Fig. S1. To enable a qualitative comparison of the denoised images and the raw images, the minimum displayed value for each image was set to the 5*^th^* percentile of the image intensity distribution.

##### Tracing single cells: correction for gastruloid rotation and drift

To make the three-dimensional time lapse data comparable to the two-dimensional simulated data, maximum z-projections of the denoised image stacks were generated for each time point using Fiji. In total, 14 mCherry-GPI-expressing cells were manually traced in Fiji using the point tool. From these 14 cells, 4 cells were dividing during the time lapse. For those cells, both daughter cells were traced after division. To compare the trajectories of these 14 cells to trajectories of cells moving in a simulated elongating gastruloid, we also traced simulated cells *in silico*. For visualization, we colored 20 of the 200 cells grey (see Video 2) and stored their ccenter positions separately. Eventually, trajectories of 12 cells were selected for the plot to show representative trajectories while minimizing the overlap between them. Before the trajectories of both *in vitro* and *in silico* cells were further analyzed or plotted on the images, cell positions were corrected for shifts due to changes in the gastruloid’s orientation over time (correction for rotation), and due to changes in the gastruloid’s center position over time (correction for drift). To calculate and correct for these shifts, we first transformed the images of the time lapse or simulation into a mask using tools from the *OpenCV* Python package (28). For the time lapse images, we first applied a Gaussian blur (kernel = 51 *×* 51 pixels) to the images. Next, we applied a threshold of 40/255 to create a binary image. Due to a slight difference in overall image intensity at *t = 1* and at *t = 26* with respect to the other images, we used a different threshold for these time points (63/255 and 45/255, respectively). To fill the holes in the resulting binary image, we applied dilation (kernel = 5 *×* 5 pixels, iterations = 40), followed by erosion (kernel = 5 *×* 5 pixels, iterations = 40). For the simulated images, we transformed the already binary images into a mask by applying erosion (kernel = 5 *×* 5 pixels, iterations = 4) followed by dilation (kernel = 5 *×* 5 pixels, iterations = 4). Next, we fitted an ellipse to the masks using the EllipseModel function from the scikit-image Python package (29). From the fitted ellipse, we obtained the center coordinates and the angle *θ* between the long axis of the ellipse and the nearest orthogonal axis when rotating counter-clockwise (x-axis for the *in silico* gastruloid, y-axis for the *in vitro* gastruloid). Often, the EllipseModel function reported a *θ* that was *π/*2 off with respect to the actual *θ*. We detected and corrected *θ* for this error by comparing *θ* with *θ* from neighboring time points or Monte Carlo steps. Finally, to correct for changes in *θ* over time due to rotation of the gastruloid, we applied a rotation matrix to the cell positions, which rotates the long axis of the ellipse an amount of *θ* to the nearest orthogonal axis (see Fig. S2A). To correct for the gastruloid’s drift over time, we corrected the cell positions for changes in the center coordinates of the fitted ellipse found for each time point or Monte Carlo step. To plot the corrected trajectories on the final figure of the time lapse or simulation, we rotated the trajectories such that they would match the gastruloid’s final orientation. For the *in vitro* gastruloid, the initial and final time points were 91 h and 96 h, respectively (Fig. 5A). For the simulation, we chose 20,000 Monte Carlo steps (MCS) as the starting point, since the elongation axis of the gastruloid was not well-defined before. For the final time point, we used 100,000 MCS. In Fig. 5C, we plotted each trajectory at 17 time points equally distributed between 20,000 and 100,000.

##### Time lapse: changing-shape analysis

To visualize the gastruloid’s changing shape during the time lapse, polygons were drawn on top of the maximum z-projection of the gastruloid at the first and last time point of the time lapse, connecting the same cells (Fig. 5B). To further analyze if the 2D shape showed lengthening and narrowing as expected from CE, we compared the length of the long and short axes of the fitted ellipses (see previous section) between the first and final time point. We also looked at the increase in overall size by comparing the size of the 2D areas obtained from the binary images.

##### Analysis of cell movement with respect to the long and short axis

We calculated the distance a cell travelled along the long axis of the gastruloid during the time lapse (91 h - 96 h) or simulation (20,000 - 100,000 MCS) (y-axis in Fig. 5E) and plotted this distance against the position of each cell at the final time point or Monte Carlo step projected on the long axis (x-axis in Fig. 5E). See Fig. 5D for a schematic illustration. The distance travelled along the short axis of the gastruloid during the time lapse or simulation (y-axis in Fig. S2B) was calculated in a similar way (see Fig. S2A), bottom panel for a schematic illustration). We also considered the direction a cell travelled along the short axis of the gastruloid during the time lapse or simulation and plotted the travelled distance along the short axis (y-axis in Fig. S2C, bottom panel) against the position of each cell at the final time point or Monte Carlo step, projected on the long axis (x-axis in Fig. S2C, bottom panel). To visualize the direction of movement along the short axis, we colored the cells by moving inwards or outwards along the short axis. Cells moving inwards (towards the short axis) were colored orange, cells moving outwards (away from the short axis) were colored dark blue, and cells that moved outwards but crossed the short axis during the time lapse or simulation were colored light blue (see Fig. S2C, top panel).

To visualize, for each cell in the simulated gastruloid, which direction it had moved, we plotted the net movement (final position - initial position) as a vector on top of each cell’s center position at the final time point. To calculate this net movement, we used the cell positions that were corrected for rotation and drift as described in the previous paragraph. For visualization, the size of the vector was divided by a factor of 20 to avoid overlapping arrows (Fig. 5F).

#### Simulations

##### Cellular Potts model

Simulations were performed using the Cellular Potts model, as first introduced in (16; 17). The Cellular Potts model simulates cells on a regular lattice Λ *⊂* ℤ^2^ and represents cells as patches of interconnected lattice sites. Each lattice site 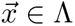 is associated with a spin, or *cell ID*, 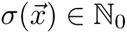. A cell *s* is then defined as the set of, usually connected, lattice sites of spin *s*, 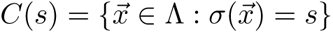. Spin *σ* = 0 is a special state reserved for the medium *M*.

The state of the system is given by a Hamiltonian, *H*, that describes the balance of forces between all cells *C* in the system. The Hamiltonian is minimized using Metropolis dynamics, by making random attempts to extend one domain *C*(*s*) by one lattice site at the expense of an adjacent domain. More precisely, during one of these copy attempts, a pair of adjacent lattice sites, 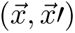 is selected at random, with 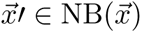 and 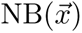 the Moore neighborhood of 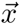, i.e., the set of directly and diagonally adjacent sites to 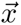. Δ*H* then becomes the energy change resulting from a copy the spin of 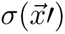 to 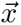. The probability that this change is accepted is assumed to follow a Boltzmann distribution with temperature *T* :

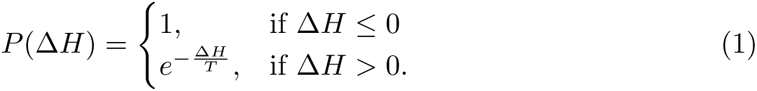

In this model, *T* determines cell motility. The copy attempts are meant to mimic pseudopodal extensions and therefore represent physically plausible microscopic dynamics. Consequently, we can consider both the equilibrium state of the system and its temporal dynamics. Time proceeds in units of Monte Carlo steps (MCS), where one step is defined as *|*Λ*|* copy attempts, i.e., the number of lattice sites in the lattice in a classical Cellular Potts implementation.

The general Cellular Potts model is given by the Hamiltonian,

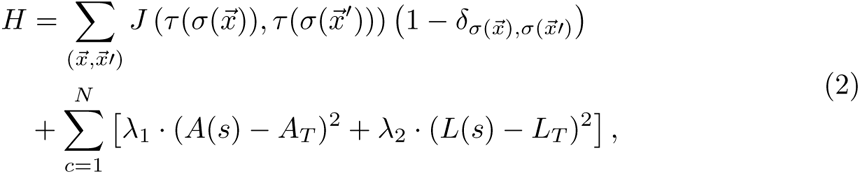

where the first sum is over all pairs of adjacent lattice sites and the second sum is over all cells. *A_T_* and *L_T_* are a target area and a target length. 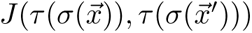 is the interfacial energy per unit length between cells, where *τ* ∈ M, 1, 2, displayed as white, red and yellow, respectively, denotes a cell type, and *δ* is the Kronecker delta. This Hamiltonian is extended with additional terms in each of the models, as defined below.

We further define surface tensions as (17),

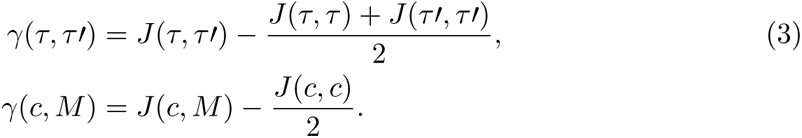

If *γ*(*c, M*) *>* 0, cells tend to stay connected, whereas if *γ*(*c, M*) *<* 0, cells tend to disperse. Similarly, if *γ*(1, 2) *>* 0, cell types 1 and 2 tend to sort, whereas cell types tend to mix if *γ*(1, 2) *<* 0.

Each simulation used a different random seed. Unless stated otherwise, we carried out 100 independent simulations for each set of parameters.

##### Differential adhesion model

The adhesion gradient was implemented by defining a gradient of 10 cell types *τ* ∈ {1, …, 10}. The cells had an adhesion energy of 25 with respect to the medium, and adhesion between cell types *τ* and *τ'* was given by *J* (*τ, τ′*) = *O* + *|τ − τ′| · S.* Here *O* indicates the offset and *S* indicates the slope for the different adhesion energies. Therefore, we obtained:

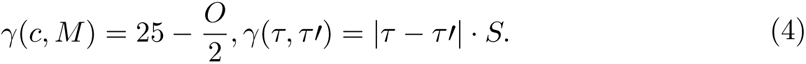

During a simulation, *O* and *S* were fixed. We achieved the different gradients we studied by changing *O* and *S* between simulations.

##### Filopodial-Tension model

In order to model CE, we added an extra term to the Hamiltonian, based on the Filopodial-Tension model proposed in (19). In this model, cells that are close to each other, but not necessarily adjacent, can exert a pulling force on each other by means of filopodia oriented in parallel to a polarization *P* (*σ*) assigned to each cell *C*(*σ*). From the cell centers, we created a set of up to *n*_max_ filopodia at random within a cone of angle *θ*_max_ and radius *r*_max_ along the long axis of the cell and in two opposing directions away from the cell (Fig. 3A). These filopodia can connect to adjacent cells’ centers of mass if these are within the cone. Filopodia are refreshed after *t*_interval_ Monte Carlo steps. The pulling force exerted by the filopodia is given by an additional term in Δ*H*,

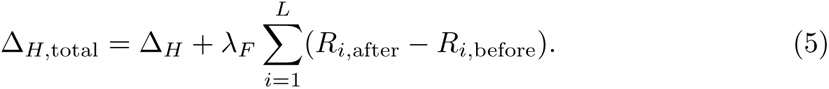

for some constant *λ_F_*. We take the sum over all filopodia, where *R_i,_*_before_ is their current length and *R_i,_*_after_ their new length resulting from the copy attempt.

Every time filopodia are refreshed, the cells’ polarization is adjusted (Fig. 3A). Consider a cell *C*(*σ*) with polarization direction *P_σ_* that has attached filopodia to cells *C*(*σ*_1_)*, …, C*(*σ_n_*) with polarization directions *P_σ_*_1_ *, …, P_σn_*. The average polarization of the neighbors is then

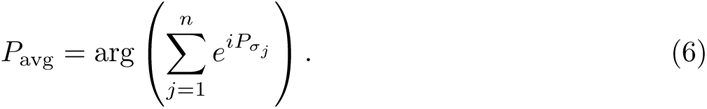

and the new polarization of cell *x* is computed by

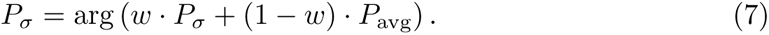

In our simulations, we used *w* = 0.99, but *w* = 0.85 was chosen in Fig. 3A for illustrative purposes. Although this polarization adjustment was already suggested by (19), they still assumed that there was a preferred direction to begin with. We have removed this assumption altogether: All cells started with a uniformly random polarization direction in the range [0, 2*π*).

##### Default parameters

A list of default parameters can be found in Table 1. These were only changed when explicitly mentioned in the manuscript.

**Table 1.** Default parameters used in the simulations.

| Symbol | Meaning | Default value |
| --- | --- | --- |
|  | Repetitions per parameter set | 100 |
| $T$ | Temperature | 50 |
| $A_T$ | Target area | 100 |
| $L_T$ | Target length | 10 |
| $\lambda_1$ | Area constraint strength | 5 |
| $\lambda_2$ | Length constraint strength | 5 |
| MCS | Number of Monte Carlo steps | 100,000 |
| $N$ | Number of cells | 200 |
| $n_{NB}$ | Number of neighbors per pixel | 8 (Moore) |
| $\theta_{\max}$ | Maximal angle for filopodia | 45 |
| $r_{\max}$ | Length of cone in target cell lengths | 2 |
| $n_{\max}$ | Maximum number of filopodia per cell | 3 |
| $t_{\text{interval}}$ | Number of MCS filopodia stay attached | 20 |
| $w$ | Memory parameter for polarization | 0.99 |
| $J(c, M)$ | Interaction energy with medium | 10 |
| $J(c, c)$ | Interaction energy between cells of the same type | 10 |

##### Implementation

We have implemented the CPM using Tissue Simulation Toolkit (30; 31). To improve the speed of simulations in Tissue Simulation Toolkit we implemented an *edgelist algorithm*, which selects only edges, *i.e.* pairs of adjacent lattice sites 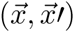 with unequal spins 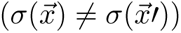. Thus, the edgelist algorithm keeps track of those pairs of lattice sites that change the configuration if copied, leading to improved efficiency especially for simulations with large cells.

Let an edge *e* be defined by a pair of adjacent lattice sites, 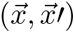 with 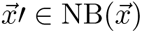, and *E* the set of edges *e* at the cellular interfaces, i.e., 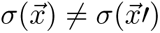. The edgelist algorithm assigns to each edge *e* ∈ *E* a unique integer from 0 up to *|*Λ*| · n_NB_ −* 1, where *|*Λ*|* is the lattice size and *n_NB_* the number of neighbors per lattice point. During initialization, the code constructs two coupled lists, called `edgelist` and `orderedgelist` to store all edges in *E*. In the list `edgelist` all edges are stored as a flattened 3D array, such that the outermost loop runs over the y-coordinate, the middle loop runs over the x-coordinate and the innermost loops runs over the neighbors of a lattice point, where the boundary state of the lattice is excluded. I.e., the *i*’th entry of the list corresponds to neighbor *nb* = (*i* mod *n_NB_*) + 1 of lattice point *p* = *Li/n_NB_J*, which can be found at position 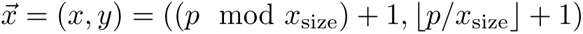. If 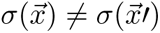 for lattice site 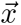 and its *nb*’th neighbor 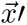, `orderedgelist` will receive a unique number from 0 to *|E| −* 1 at position *i* and a value of *−*1 otherwise. In `orderedgelist`, the first *|E|* entries indicate the locations of the entries of `edgelist` with values larger than *−*1, while the subsequent entries are equal to *−*1 (Table 2). This assures a 1-to-1 relationship between the values in one list and the positions with values unequal to *−*1 in the other list.

**Table 2.**
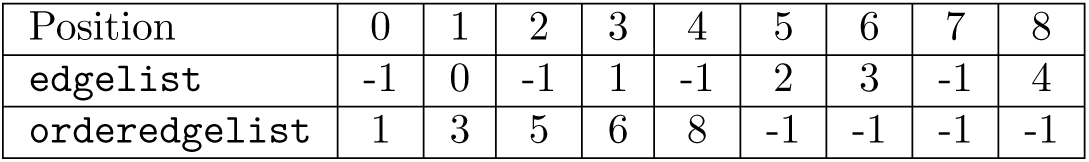
Example of edgelist arrays

For every copy attempt, i.e., an attempt to copy a lattice site to an adjacent lattice site, we sample a random edge from `orderedgelist`. If a copy attempt is accepted, both lists are updated accordingly by iterating over the neighborhood of the changed pixel. New edges are added at position *|E|* in `orderedgelist`. If an edge at position *i* in `orderedgelist` has to be removed, the edge at the last position *|E| −* 1 will replace this edge such that the `orderedgelist` remains a consecutive list. The list `edgelist` is also updated such that a 1-to-1 relationship between the two lists is preserved. This relationship is essential to efficient updating and sampling from these lists. Furthermore, the number of copy attempts per Monte Carlo step was changed to *|E|/n_NB_* copy attempts per Monte Carlo step, where *|E|* may vary during a Monte Carlo step, from the *|*Λ*|* copy attempts in the classical algorithm. This rescaling is justified because a copy attempt in the classical Cellular Potts model has a probability of 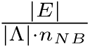to select a pair of lattice sites belonging to two different cells and hence *|E|/n_NB_* of such pairs are selected per classical Monte Carlo step on average.

In order to schedule simulations efficiently, we have used GNU parallel (32) on the high performance cluster ALICE provided by Leiden University.

#### Statistics

##### 2D Kolmogorov-Smirnov test

We used the two dimensional version of the Kolmogorov-Smirnov test (Fasano-Franceschini test) (33) to determine which model variant and which parameter set gave the best fit to experimental data. The test statistic *D* gives a measure of similarity between two distributions. Two identical distributions will give *D* = 0, whereas two fully separated distributions of data points result in *D* = 1. Fully separated here means that we can find a point (*x, y*) such that the two distributions are in opposite quadrants when drawing axes through (*x, y*). A *p*-value was computed for the null hypothesis that the two empirical distributions were drawn from the same underlying distribution. Within a model, the best parameter set was considered the one giving the lowest value of *D*, and the highest *p*-value, comparing 100 simulations to the full experimental data set. Similarly, the best model variant (in the case of two cell types) was selected based on *D* for the respective optimal parameter sets. The Fasano-Franceschini test was calculated using an existing implementation in R (34).

#### Code availability

Code for our simulations and LOCO-EFA can be found on GitHub: https://github.com/Martijnadj/Gastruloids_convergent_extension_differential_adhesion

## Results

### Gastruloid elongation leads to a wide variety of shapes

We generated gastruloids according to the original protocol developed by (4) and visualized their shapes by wide-field fluorescence imaging at multiple time points. After 48 h the cells had formed spherical aggregates (Fig. 1A). At this time point the Wnt pathway is activated by adding a GSK-3 inhibitor. Among many other functions, the Wnt pathway regulates AP axis formation. Consequently, gastruloids started to elongate after approximately 72 h leading to a variety of shapes by 96 h (Fig. 1A). To enable quantification, we extracted shape outlines from the wide-field images, which are two-dimensional projections of the (three-dimensional) gastruloid shapes. To that end, we developed a computational tool that segments gastruloids and represents their shapes as individual binary masks (Fig. 1B), see Materials and Methods). We analyzed the shapes with Lobe-Contribution Elliptic Fourier Analysis (LOCO-EFA) (15), which is more easily interpretable than conventional elliptic Fourier analysis. LOCO-EFA decomposes a shape outline into a series of modes, such that each mode has a defined number of lobes. A coefficient *L_n_* then quantifies the contribution of the mode with *n* lobes to the shape. Thus, while the first component (*L*_1_) indicates the overall (linear) size of a shape, *L*_2_ and *L*_3_ indicate the presence of 2 or 3 lobes, respectively. Larger values of *L*_3_*/L*_1_ thus indicate more complex shapes, which are not just simple ellipsoids. LOCO-EFA analysis revealed that gastruloids exhibit more complex shapes more often at 96 h than at 72 h (Fig. 1C). Modes with more than three lobes had smaller contributions and are more sensitive to the noise introduced by the stochastic simulation algorithm. Therefore, we restricted our analysis to *L*_2_ and *L*_3_. All in all, our analysis showed that gastruloids assume shapes that are more complex than simple ellipsoids.

### Differential adhesion is insufficient to explain the observed shapes

We next wanted to find the simplest mechanistic explanation of the observed shape distribution. First, we argued that a homogeneous population of cells will not produce elongated shapes, even in the presence of cell migration and proliferation. Due to the adhesion between cells and the medium, a homogeneous cell aggregate will always be approximately spherical, as that shape minimizes cell interactions with the medium at fixed total volume. The cell aggregate thus effectively behaves like a liquid droplet with positive surface tension. Gastruloids are, however, heterogeneous and contain multiple cell types at 96 h (4; 35). These cell types likely differ in their adhesive properties. The simplest hypothesis to explain formation of elongated shapes might thus be cell-to-cell differences in the strength of adhesion, as first explored in the CPM-based model of differential-adhesion-driven convergent-extension by (36). This model is parameterized by *γ*(*c, M*), the surface tension between any cell *c* and the medium *M*, and *γ*(*τ, τ′*), the surface tension between cell type *τ* and cell type *τ′* (see Materials and Methods). The model assumes a linear gradient of adhesion differences *γ*(*τ, τ′*) = *S · |τ − τ′|*, where *S* is the slope of the gradient. In our simulations of the differential adhesion model we assumed 10 different cell types, which were initially arranged according to their adhesive properties. This initial arrangement is consistent with the occurrence of symmetry-breaking already prior to elongation in gastruloids (4). We confirmed the presence of multiple cell types in our gastruloid experiments by staining of Brachyury, a marker of primitive streak-like cells. We observed these cells to be strongly localized after gastruloid elongation (Fig. 2A).

**Fig 2.**
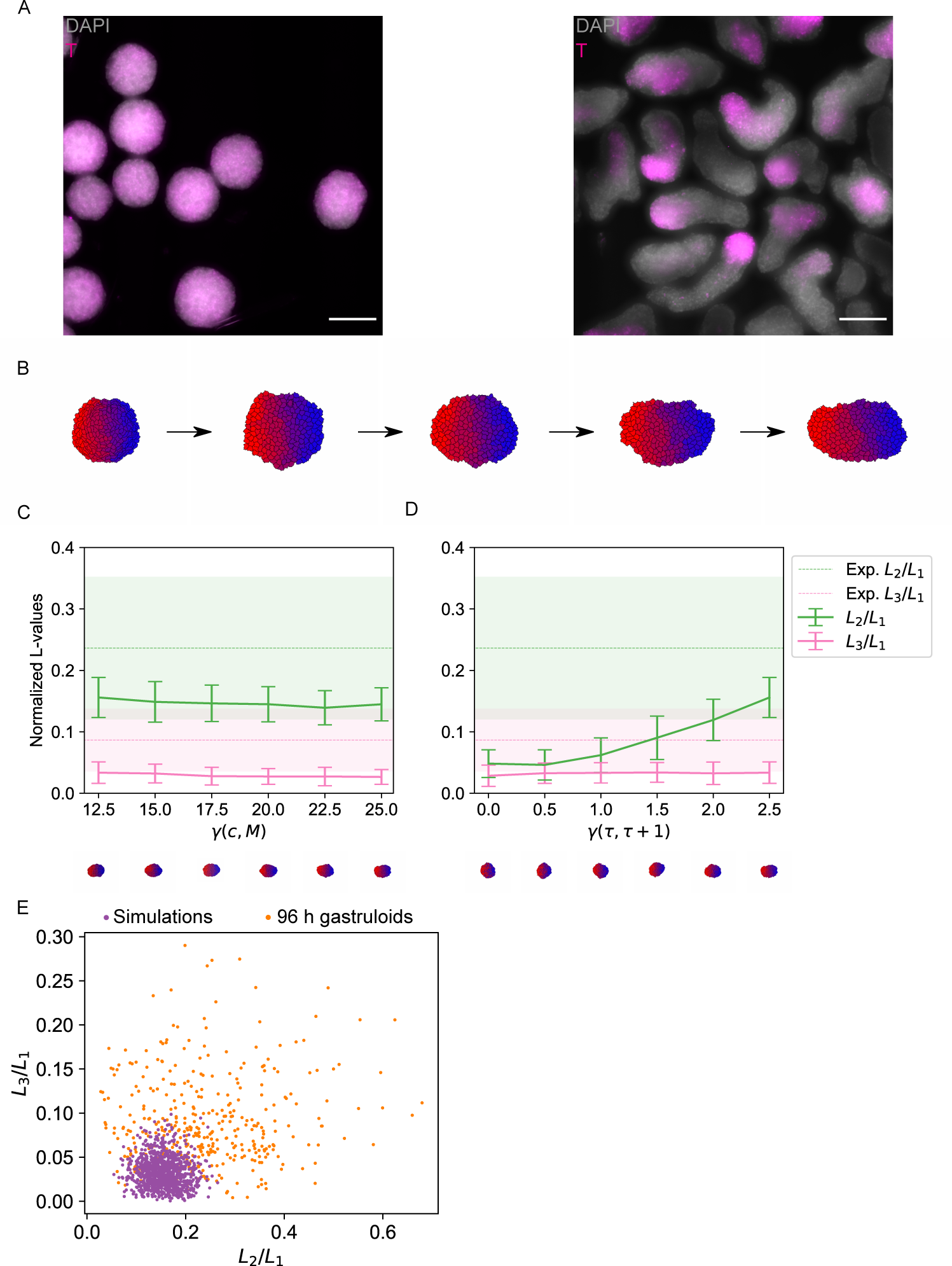
Differential adhesion cannot explain the shape distribution of gastruloids. A: Wholemount immunostaining of Brachyury in 72 h and 96 h gastruloids. Shown is a maximum z-projection. Cell nuclei were stained with DAPI. Scale bars: 200 *µ*m. B: Example shapes resulting from a simulation of differential adhesion for 120, 30,000, 60,000, 90,000 and 120,000 Monte Carlo steps. Different cell types / adhesion strengths are indicated by color. C,D: Quantification of shapes resulting from simulations of differential adhesion. Shown are LOCO-EFA coefficients averaged over a population of shapes. The shaded areas (in the experimental data) and vertical bars (in the simulation data) indicate standard deviations. C: The surface tension *γ*(*c, M*) was varied. The slope was kept constant at a value corresponding to a tension *γ*(*τ, τ* + 1) between two adjacent cell types of 2.5. D: The slope of the adhesion gradient was varied. The offset was kept constant at 25. *L*_2_*/L*_1_ increases with increasing slope but *L*_3_*/L*_1_ remains approximately constant. E: Scatter plot of the scaled LOCO-EFA coefficients *L*_3_*/L*_1_ versus *L*_2_*/L*_1_ for the experimental data and simulations of differential adhesion with offset 25 and slope 2.5

As hypothesized, the differential adhesion model produced elongated shapes after 120,000 Monte Carlo steps (Fig. 2B). However, more detailed analysis revealed that this model is not able to explain the experimentally measured shapes. We ran multiple simulations with different values for two parameters: the baseline interfacial tension of the gastruloid surface *γ*(*c, M*) and the slope *S* of the adhesion gradient. Larger values of *γ*(*c, M*), which correspond to higher surface tension, resulted in slightly smaller LOCO-EFA coefficients for 2 or 3 lobes (Fig. 2C). In other words, higher surface tension results in smaller deviations from a circular shape, as to be expected. Increasing the slope parameter *S* increases the interfacial tension *γ*(*τ, τ* + 1) between neighboring cell types *τ* and *τ* + 1. A higher *γ*(*τ, τ* + 1) reduces the mixing of cell types and should thereby increase elongation. As expected, we observed an increase in *L*_2_*/L*_1_ with increasing *S* (Fig. 2D). The value of *L*_3_*/L*_1_, however, stayed approximately constant. Since the tissue started to disintegrate in simulations with slopes higher than the ones shown, we were unable to obtain values of *L*_2_*/L*_1_ or *L*_3_*/L*_1_ observed in experiments (Fig. 2E). Differential adhesion thus does lead to somewhat elongated shapes, but elongation is less pronounced than observed in experiments. Differential adhesion also does not seem to drive formation of shapes with more than two lobes.

### A filopodial-tension model with self-organized mediolateral axis recapitulates the experimental results

Having found that differential adhesion is insufficient to explain the observed shapes, we adapted the model of CE by Belmonte et al. (19) for our simulations. This model is based on the “crawling” model of CE. It assumes that cells exert pulling forces upon one another by filopodia-like protrusions extended preferentially perpendicular to a pre-defined polarization axis. These protrusions are transient and new protrusions are extended periodically. Due to the pulling, the tissue elongates along the pre-defined axis and narrows in the perpendicular direction, as expected in CE.

There is no reason to assume that there is an extrinsically imposed mediolateral orientation in gastruloids. Therefore, we modified the model by Belmonte et al. (19) such that the elongation axis emerges in a self-organized way. At the start of the simulation we assigned a uniformly random polarization to each cell. At each Monte Carlo step a new polarization is determined from a weighted average of a cell’s polarization and the polarizations of all neighbors connected through protrusions (Fig. 3A). The weight on the cell’s own polarization is chosen to be high (typically around 0.99), which reflects a tendency of the cell to preserve its polarization over time. Due to this mechanism, cells become locally aligned, allowing CE to take place (Fig. 3B).

**Fig 3.**
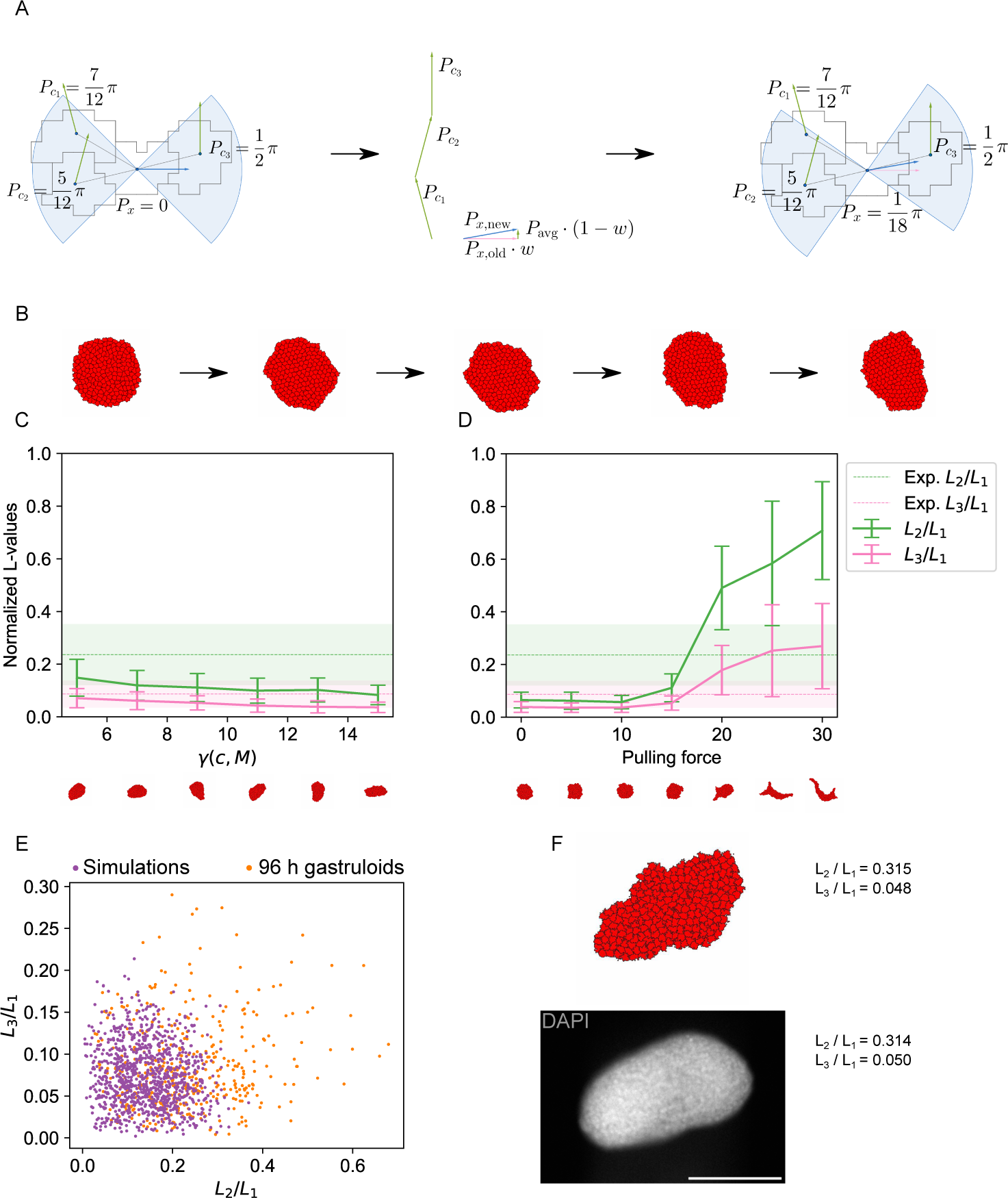
Simulations including CE reproduce average LOCO-EFA coefficients. A: Schematic of the filopodial tension model with self-organized preferred direction. The new polarization direction of a cell is the weighted average of its old polarization direction and the polarization directions of all neighboring cells connected by filopodia. B: Example shapes resulting from a simulation with one cell type for 100, 25,000, 50,000, 75,000 and 100,000 Monte Carlo steps. C,D: Quantification of shapes resulting from simulations of CE. Shown are LOCO-EFA coefficients averaged over a population of shapes. The shaded areas (in the experimental data) and vertical bars (in the simulation data) indicate standard deviations. C: The interaction strength with the medium was varied. The pulling force was kept constant at 15. D: The pulling force was varied. The surface tension with the medium was kept constant at 5. E: Scatter plot of the scaled LOCO-EFA coefficients *L*_3_*/L*_1_ versus *L*_2_*/L*_1_ for the experimental data and simulations of CE with pulling force *λ_F_* = 15 and surface tension *γ*(*c, M*) = 5. These parameters gave the smallest 2D Kolmogorov-Smirnov test statistic *D* = 0.40 and *p* = 9.3 . 10*^−^*^9^. F: Example of experimentally observed shape (bottom) with highly similar simulated shape (top) and their corresponding scaled LOCO-EFA coefficients *L*_3_*/L*_1_ and *L*_2_*/L*_1_. Bottom: a 96 h fixed gastruloid. Shown is the mid-plane of a z-stack. Cell nuclei were stained with DAPI. Scale bar: 200 *µ*m.

The shapes that we obtained with this new model depended most strongly on two parameters: the surface tension between medium and cells, *γ*(*c, M*), and the pulling force *λ_F_*. Increasing the surface tension *γ*(*c, M*) resulted in more circular shapes (Fig. 3C), as such shapes reduce the length of the costly interface between cells and medium. We also observed that increasing the pulling force *λ_F_*, produced less circular shapes (Fig. 3D). Using a simple grid search we found parameters that resulted in average LOCO-EFA coefficients similar to the experimentally observed values. Notably, for a pulling force between 15 and 20, both *L*_2_*/L*_1_ and *L*_3_*/L*_1_ match the experimental values on average with the surface tension kept constant at 5. A simultaneous match was not found for the surface tension within our parameter range.

Average LOCO-EFA coefficients are of course an incomplete description of the shape distributions, in particular if these distributions are not unimodal. Hence, we wanted to compare experimental and simulated shape distributions directly. For this comparison to be meaningful, the variability among multiple simulation runs should not just be due to the inherent stochastic nature of the Monte Carlo algorithm, but reflect the energetic landscape of gastruloid shapes. To explore the nature of the fluctuation in our simulations we ran simulations of extended length and visualized their trajectories in LOCO-EFA coefficient space (Fig. S3). After an initial phase of fast movement, the simulation trajectories kept exploring a fairly large area in coefficient space. This supports the notion that the observed variability is not just given by stochastic fluctuations around a sharp global energy minimum but rather reflects a (locally) flat energy landscape. This energy profile underlies both the variability among simulations and the variety of gastruloid shapes. It is thus meaningful to compare the simulated and measured shape distributions.

To compare distributions quantitatively, we used a two-dimensional Kolmogorov-Smirnov (KS) test on the joint distribution of the LOCO-EFA coefficients. The null hypothesis of this test assumes that two empirical samples are drawn from the same underlying distribution. The corresponding test statistic *D*, which lies between 0 and 1, is small if the two empirical distributions are similar and close to 1 if they are very distinct. We also calculated a p-value for the rejection of the null hypothesis. We considered those model parameters to be optimal, for which we found the smallest test statistic *D* when comparing experimental and simulated shape distributions. For our CE model, optimal parameters (pulling force = 15 and *γ*(*c, M*) = 5) led to average shape coefficients similar to those observed experimentally (Fig. 3C,D) but the distributions remained distinct (*D* = 0.40 and *p* = 9.3 *·* 10*^−^*^9^) (Fig. 3E). This indicates that the model could not fully recapitulate the experiments in terms of shape variety. Nevertheless, we were able to find several computationally created shapes that closely resembled individual experimental shapes (Fig. 3F). This demonstrates that our simulations can produce realistic shapes. All in all, the CE model produced shapes that were more similar to observed shapes compared to the differential adhesion model, but the shape distribution still differed significantly from the experimental findings. In particular, larger values of *L*_2_*/L*_1_ and *L*_3_*/L*_1_, found in the *in vitro* gastruloids, were not reproduced by simulations.

### Synergistic CE and differential adhesion model has improved agreement with experiments

Naturally, we were wondering whether a combination of CE and differential adhesion could improve the agreement with the experimental results further. To create a minimal model of cell type diversity in the context of our CE model, we added a second cell type (shown in yellow). To achieve spatial segregation of cell types, we assumed that adhesion between like cells was stronger than between cells of different types (Fig. 4A). For comparison, we also considered the case where it is energetically favorable for cell types to mix. Multiple variants of pulling are possible when there are two cell types. We investigated four different variants: (1) cells can only attach filopodia to cells of the same cell type (”same type” model), (2) all cells can attach filopodia to all other cells (”all-all” model), (3) only yellow cells can attach filopodia to other cells (”yellow-all” model) and (4) yellow cells can only attach filopodia to yellow cells (”yellow-yellow” model). Using again the 2D KS test to compare simulated and experimental shapes, we found the “same type” model to provide the best fit with the data (Fig. 4). (See Fig. S4, Fig. S5 and Fig. S6 for results obtained with the other model variants).

**Fig 4.**
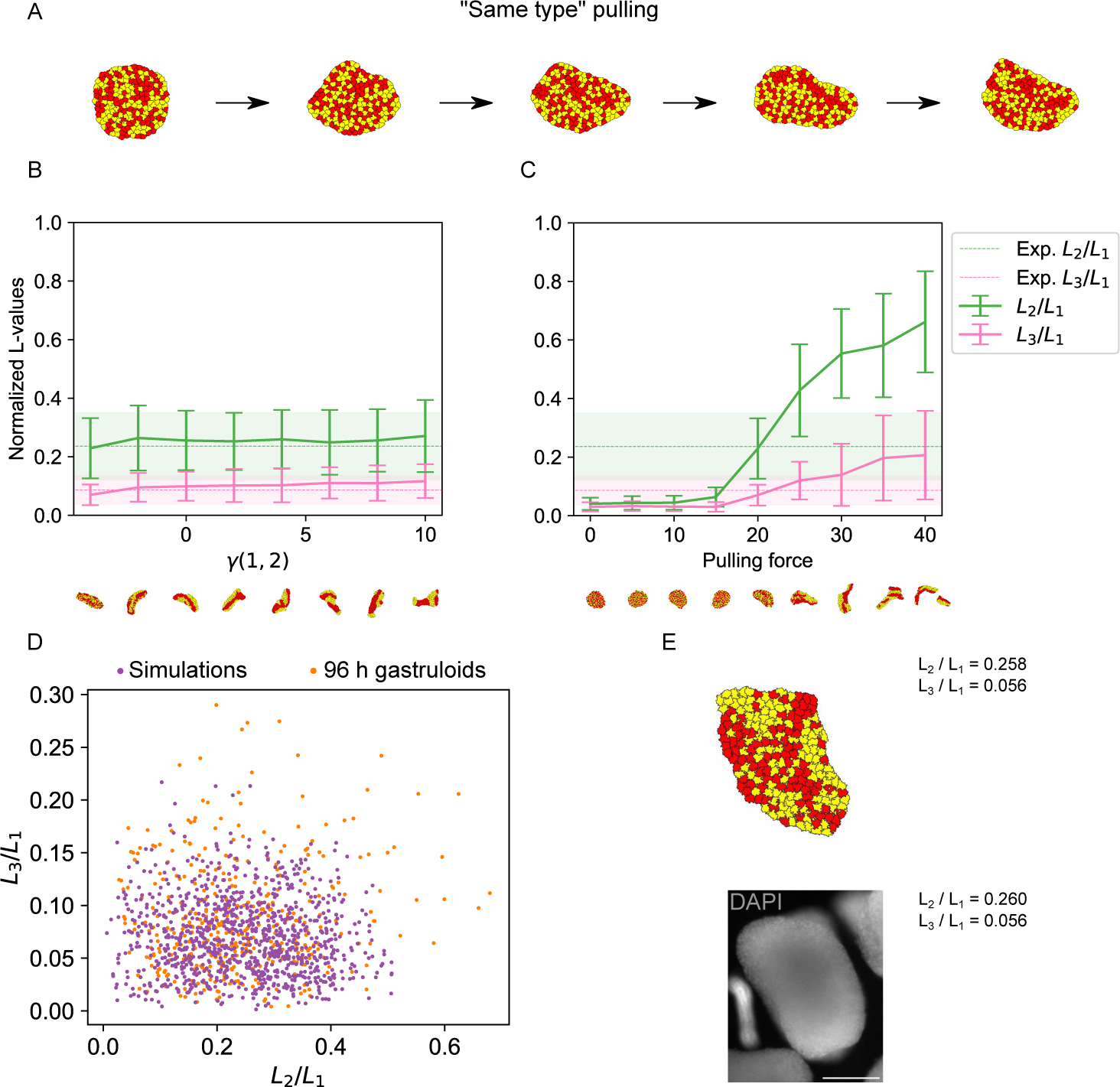
Simulations of CE with two cell types create more extreme shapes. A: Example shapes resulting from a simulation of yellow-all pulling for 100, 25,000, 50,000, 75,000 and 100,000 Monte Carlo steps. B,C: Quantification of shapes resulting from simulations of CE with two cell types. Shown are LOCO-EFA coefficients averaged over a population of shapes. The shaded areas (in the experimental data) and vertical bars (in the simulation data) indicate standard deviations. B: The surface tension between the two cell types was varied and the pulling force was kept constant at 20. C: The pulling force was varied. The surface tension between the two cell types was kept constant at −4. D: Scatter plot of the scaled LOCO-EFA coefficients *L*_3_*/L*_1_ versus *L*_2_*/L*_1_ for the experimental data and simulations of CE with pulling force *λ_F_* = 20 and surface tension *γ*(1, 2) = *−*4. These parameters gave the smallest 2D Kolmogorov-Smirnov test statistic *D* = 0.15 and largest p-value *p* = 0.15. E: Example of experimentally observed shape (bottom) with highly similar simulated shape (top) and their corresponding scaled LOCO-EFA coefficients *L*_3_*/L*_1_ and *L*_2_*/L*_1_. Bottom: a 96 h fixed gastruloid. Shown is the mid-plane of a z-stack. Cell nuclei were stained with DAPI. Scale bar: 200 *µ*m.

Both *L*_2_*/L*_1_ and *L*_3_*/L*_1_ remained almost constant when the surface tension *γ*(1, 2) between cell types was increased (Fig. 4B). Increasing the the pulling force, however, led to a strong increase in the average values of *L*_2_*/L*_1_ and *L*_3_*/L*_1_ (Fig. 4C). Simulated average LOCO-EFA coefficients were similar to the experimental results at a pulling force between 15 and 20 with the surface tension between the two cell types kept constant at −4. A simulation with a single set of fixed parameters is thus able to reproduce the average experimental shape. Note that a negative surface tension causes cell types to be mixed as it favors a larger interface between them. To make a more detailed comparison we considered the complete distribution of shapes and used the 2D KS test (Fig. 4D). For a pulling force of 15 and surface tension of −4, we found a test-statistic of *D* = 0.15, which is much lower than for the model with one cell type. Correspondingly, the p-value was high (*p* = 0.15), which means that there is no indication of simulated and experimental results being different, given the available data. Simulated gastruloids with two cell types exhibited slightly higher values of *L*_2_*/L*_1_ and *L*_3_*/L*_1_, compared to homogeneous gastruloids (Fig. 3E). One explanation for this effect could be that clusters of the same cell type tend to become circular, which leads to slightly more complex shapes. Due to the pulling of cells of the same cell type, the cells still sorted slightly, even with a negative surface tension. Once again, we found that simulations and experiments with similar LOCO-EFA coefficients resulted in similar shapes (Fig. 4E). Overall, adding another cell type improved the agreement of the CE model with the experimental observations.

The 2D KS test identified the “same type” model as the optimal model in the sense that it achieved the lowest value for the test statistic *D*. By itself, this cannot rule out the other model variants. In Fig. S7 we report the test statistic *D* for all model variants and various model parameters. Differential adhesion performed poorly for all tested parameters (Fig. S7A). In the CE model with one cell type, there is a clear correlation between surface tension *γ*(*c, M*) and pulling force values that result in good models (Fig. S7B): If the pulling force is increased, the surface tension must be increased, as well. For CE with two cell types, the relationship between optimal surface tension *γ*(1, 2) and pulling forces is reversed (Fig. S7C-F): Higher pulling forces need to be compensated with lower surface tensions between the two cell types. Notably, for a pulling force larger than 20, the test statistic dropped rapidly for the “same type”, “all-all” and “yellow-all” variants (Fig. S7C, D, F), because shapes become highly irregular in that regime. For the largest pulling forces we considered, the tissues even occasionally split into multiple disconnected parts. Furthermore, the “all-all” variant produced shapes that were significantly different (*p* = 1.4 *·* 10*^−^*^7^) from the measured shapes for all parameter values (Fig. S7D), while the difference was not significant for the other model variants after Bonferroni correction (*p* = 0.15*, p* = 0.0098 and *p* = 0.12 respectively) (Fig. S7C, E, F). In conclusion, among the model variants tested, the “same type” and “yellow-yellow” models matched the experimental data well for a range of model parameters. Direct measurements of cell adhesion or exerted forces would be necessary to conclusively decide between the model variants.

### Time lapse imaging of gastruloids reveals hallmarks of CE

We next looked for direct evidence of CE in gastruloids. To that end we created mosaic gastruloids, consisting of mostly unlabeled, wild-type mESCs and a small fraction (1:16) of mESCs with constitutively fluorescent plasma membranes (GPI membrane anchor labeled with an mCherry fluorescent protein). These mosaic gastruloids allowed us to track individual cells over time using a light-sheet fluorescence microscope (see Fig. 5A and Video 1).

**Fig 5.**
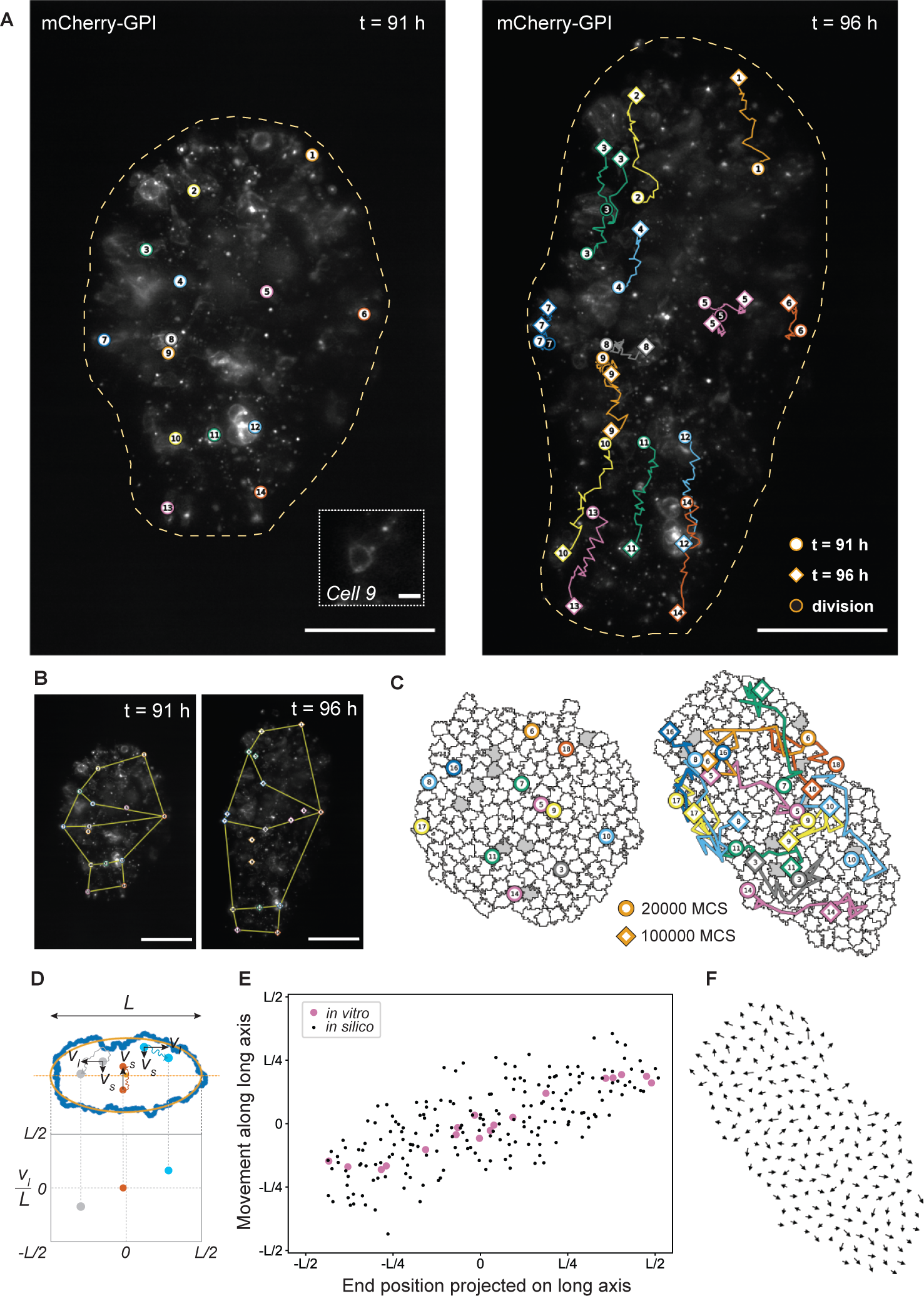
Time lapse imaging of gastruloids reveals hallmarks of CE. A: Maximum z-projection of a gastruloid after 91 h (left) and 96 h (right) of differentiation. Cell trajectories are overlayed. The outline of the gastruloid is indicated by a yellow dashed line. Approximately 1 in 16 of the cells expressed mCherry-GPI, which fluorescently labelled the membrane. 14 of these cells were traced over the duration of the experiment, which had a total of 31 time points (10 min interval). Circles with a white core show the position of the cells at 91 h. Diamond shapes show the position of the cells at 96 h. Dividing cells are indicated by a circle with a dark core. For each dividing cell, both daughter cells were traced. The inset at 91 h shows a zoom-in on cell number 9 that was traced. Scale bars: 200 *µ*m, and 10 *µ*m for the inset. B: Deformation of shapes defined by the position of selected cells. Scale bars: 200 *µ*m. C: Stills of a simulation of a mosaic gastruloid at 20,000 Monte Carlo steps (left) and 100,000 Monte Carlo steps (right), showing 20 cells in grey and 180 cells in white. Trajectories of 12 grey cells shown at 17 time points are overlayed. Trajectories were corrected for the gastruloid’s rotation and drift. Trajectories start at 20,000 Monte Carlo steps (circles) and end at 100,000 Monte Carlo steps (diamond shapes). D: Scheme showing how cell positions were projected on the long axis of a measured or simulated gastruloid. Positions of cells were rotated to have the new *x* coordinate correspond to the position on the long axis and the new *y* coordinate to the position on the short axis. The movement along the long axis *v_l_* or short axis *v_s_* was measured for each cell and normalized to the total length of the gastruloid *L*. E: Distance moved versus end position along the long axis for simulated and measured cell trajectories. Large, pink circles represent cells of the *in vitro* gastruloid. Small, black circles represent cells of the simulated gastruloid. F: Net movement (final position - initial position) for each cell plotted as a vector on top of each cell center at the final Monte Carlo step. The net movement was corrected for rotation and drift as described in the Materials and Methods. For visualization, the size of the vector was divided by a factor of 20 to avoid overlapping arrows.

A maximum z-projection of the recorded images enabled us to measure the gastruloid’s shape for each time point. We observed elongation of the gastruloid over a period of 5 h. During this period, the overall size of the gastruloid increased by approximately 38% in projected area (obtained from maximum z-projections, see Materials and Methods). This increase in size could be partially explained by cell proliferation, as some of the labeled cells divided over the course of 5 h (Fig. 5A). Using immunofluorescence staining of the cell proliferation marker Ki-67 in 96 h gastruloids we confirmed that cell proliferation did occur, but we did not observe a strong localization of proliferating cells (Fig. S8). Nevertheless, the observed elongation showed hallmarks of CE. Fitting an ellipse to the two-dimensional (2D) projected shapes revealed an increase in the long axis of 52% and a decrease in the short axis of 8% over the 5 h time period. In addition, following the deformation of simple geometric shapes defined by the positions of selected cells (Fig. 5B) we observed lengthening and narrowing of the tissue reminiscent of presomitic tissue in the mouse that undergoes CE (37). Overall, we observed cell proliferation to occur uniformly across the gastruloids and high levels of cell motility. These observations exclude elongation mechanisms that are based on strongly localized proliferation or local rearrangements in a static configuration of cells.

The time-resolved single-cell measurements also provided another opportunity to test the validity of our computational model. We ran new simulations with optimal parameters for the single cell type model obtained above (Fig. 3) and collected the positions of the cells over several thousand Monte Carlo steps, (see Fig. 5C and Video 2). To compare simulations and experiments, we projected cell positions on the long axis of the gastruloid (Fig. 5D) and plotted the distance a cell moved along this axis versus the final position of its trajectory (Fig. 5E). Both in experiments and simulations, cells ending up close to the poles of the gastruloids had travelled further and cells ending in the middle hardly moved along the long axis. This trend is consistent with CE, where we expect cells near the center of the gastruloid to move mainly inwards along the short axis (convergence). At the same time, we expect that cells further away from the center move outwards along the elongating axis of the gastruloid (extension). Projection of cell positions on the short axis did not reveal any additional trends with respect to the final position of the trajectories (Fig. S2). Interestingly, in the experimental data, cells in the first and last quarter of the gastruloid (along the long axis) seemed to move approximately by the same amount (Fig. 5E). Correspondingly, in the simulations, cells at larger distances from the short axis seemed to move towards the poles in more homogeneous groups (Fig. 5F). Taken together, the simulations recapitulated salient features of the time lapse imaging data.

### Inhibition of the ROCK pathway prevents elongation but not cell type segregation

For an additional, independent validation of the importance of CE, we considered the perturbation of signaling pathways. Prior studies in the mouse neural tube (14) and elongating aggregates of P19 mouse embryonal carcinoma cells (38) have shown that CE depends on the Rho kinase (ROCK) pathway. Growing gastruloids in the presence of a ROCK inhibitor during the final 24 h of differentiation resulted in substantially fewer elongated gastruloids and the remaining non-spherical examples often assumed a snowman-like shape (Fig. 6A).

**Fig 6.**
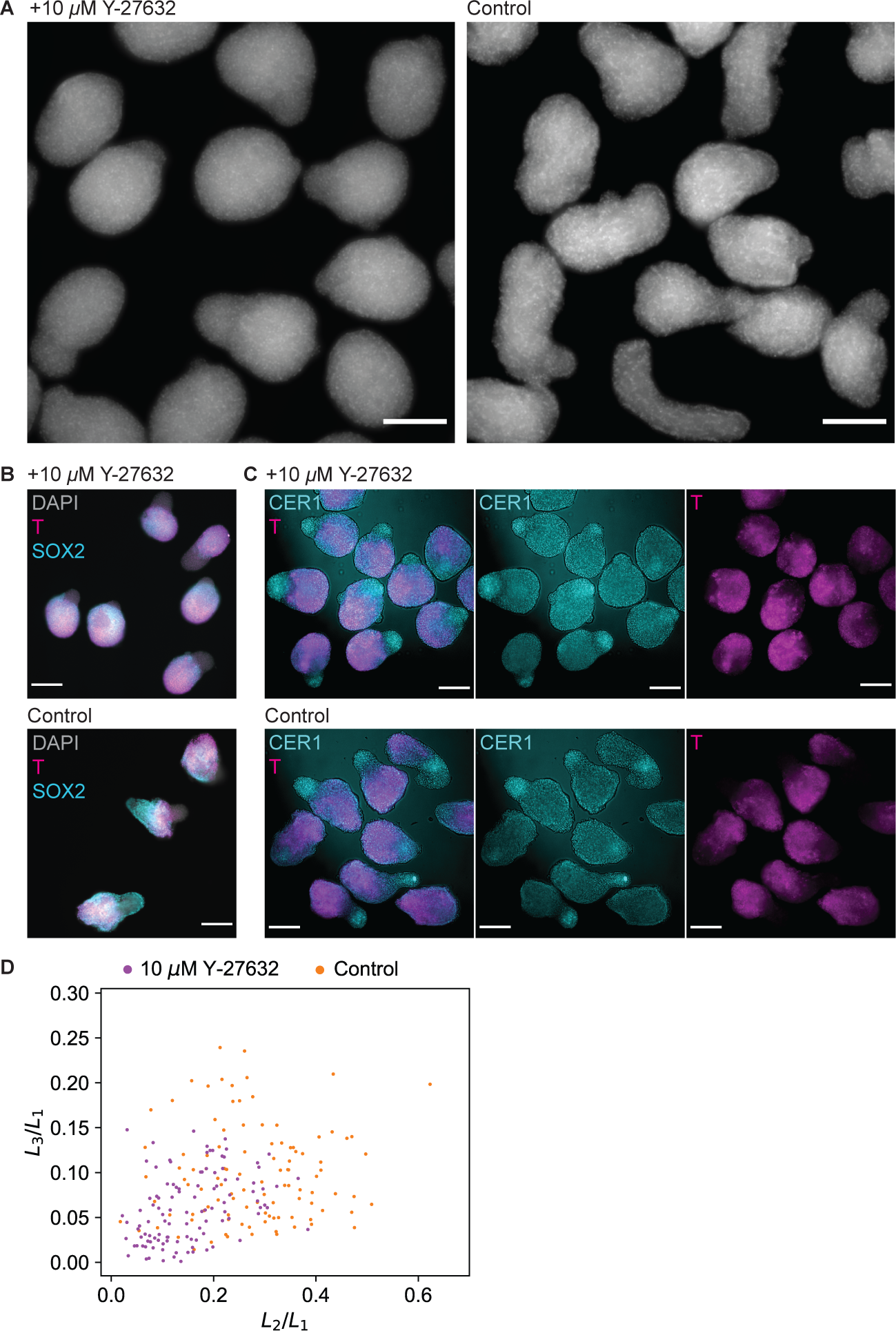
Inhibition of the ROCK pathway prevents elongation but not cell type segregation. A: Fixed gastruloids at 96 h treated with (left) or without (right) 10 *µM* of the ROCK inhibitor Y-27632 during the final 24 h of differentiation. Shown is a maximum z- projection. Cell nuclei were stained with DAPI. Scale bars: 200 *µ*m. B: Wholemount immunostaining of Brachyury/T and Sox2 in fixed gastruloids at 96 h treated with (top) or without (bottom) 10 *µM* Y-27632. Cell nuclei were stained with DAPI. Scale bars: 200 *µ*m. C: Wholemount immunostaining of Brachyury/T and Cer1 in fixed gastruloids at 96 h treated with (top) or without (bottom) 10 *µM* Y-27632. Cell nuclei were stained with DAPI. Scale bars: 200 *µ*m. D: LOCO-EFA analysis of measured gastruloids at 96 h treated with (n = 110) or without (n = 106) 10 *µM* Y-27632. Results of two biological replicates were combined. Each data point is a gastruloid. The scatter plot shows the scaled LOCO-EFA coefficients *L*_2_*/L*_1_ and *L*_3_*/L*_1_.

LOCO-EFA analysis of gastruloid shapes revealed a shift to lower scaled LOCO-EFA coefficients *L*_2_*/L*_1_ and *L*_3_*/L*_1_ as a result of ROCK inhibitor treatment (Fig. 6D). Antibody staining of markers of primitive streak (Brachyury/T), neural tissue (Sox2) or somitic tissue (Cer1 (39; 40)) revealed that the snowman shape was likely driven by segregation of different cell types (Fig. 6B-C). In control conditions, cell types were similarly separated but the gastruloids were also elongated, seemingly across multiple different cell types. These results confirm that the observed gastruloid shapes can be explained by a combination of CE - mediated by the ROCK pathway - and differential adhesion between multiple cell types.

## Discussion

In this study we showed that CE combined with differential adhesion can explain the shapes of elongating gastruloids. While differential adhesion alone was unable to produce sufficiently elongated and complex shapes on its own, CE produced, in a self-organized manner, simulated shape distributions that were more similar to the experimental results. Combining both mechanisms, we found a single set of simulation parameters that reproduced the mean and variability of experimental shapes.

We further identified that the “same type” model for two cell types resulted in a distribution of shapes that resembled the *in vitro* shape distribution best. However, we could not conclusively reject the other three models we put forward, which might be achievable when larger *in vitro* data sets become available. Alternatively, direct *in vitro* measurements of the forces between the different cell types, both with and without ROCK inhibition, could help determine whether both cell types play an active role in CE. If both cell types exert forces, the “yellow-yellow” model could be rejected, for example. The filopodial pulling mechanism we assumed in our model is primarily inspired by the “crawling” mechanism. Although we observed a large degree of cell motility, which is most consistent with “crawling”, we have insufficient data to exclude that the junction shrinking mechanism is also active. The filopodial pulling model is in fact sufficiently general to represent both mechanisms of CE, such that an explicit junction contraction model or a model combining both mechanisms (11) could yield similar gastruloid shape distributions.

In the presence of ROCK inhibitor, gastruloid elongation was inhibited, but cell types still separated, resulting in snowman-like shapes. These shapes were reminiscent of another gastruloid-based differentiation protocol that produces an endodermal compartment (41). It would be interesting to explore, whether similar cell separation mechanisms occur in both systems and whether the resulting shapes can be explained by a line tension between two fluid phases (42). Similar shapes have also been produced *in silico* in the presence of differential adhesion (36). When gastruloids were treated with ROCK inhibitor, shapes with high LOCO-EFA coefficients *L*_2_*/L*_1_ and *L*_3_*/L*_1_ disappeared. It is noteworthy that many gastruloids under control condition had similar shapes to the ROCK-inhibitor treated gastruloids. This might indicate that a significant amount of gastruloids failed to elongate properly, possibly because CE did not take place at all. It would be interesting to study whether a failure to elongate is related to different initial cell states, as suggested recently (43).

Beyond revealing a possible morphogenic process during embryonic development, the methods we used for shape quantification and simulation might also be useful in biomedical or pharmacological applications. Recently, human gastruloids have been used to test, whether particular drugs cause developmental malformations (teratogenicity) (44). In that study, shapes were analyzed with simple metrics, such as the area (of the 2D projection), aspect ratio and circularity. Here, we employed the LOCO-EFA approach, which might give a more detailed description of drug-treated gastruloids. Moreover, accompanying CPM simulations could even lead to hypotheses for a drug’s mechanism of action.

In the future, it would be very interesting, albeit computationally much more costly, to repeat the simulations in three dimensions and also incorporate cell growth and division. Furthermore, a more extensive parameter sweep with the current model could provide new insights, although this is, again, computationally expensive. The model could be further extended by including more details of the different cell types in gastruloids. On the experimental side, it would be very informative to measure the adhesion strength and forces between different cell types and resolve the changes in cell shapes during elongation. We hope that this study will serve as a stepping stone for more sophisticated models and more detailed measurements.

## Acknowledgments

This work was carried out on the Dutch national e-infrastructure with the support of SURF Cooperative and using the compute resources from the Academic Leiden Interdisciplinary Cluster Environment (ALICE) provided by Leiden University. E.A. acknowledges support by a Stichting voor Fundamenteel Onderzoek der Materie (FOM, www.nwo.nl) projectruimte grant (16PR1040). This work was also supported by NWO grants NWO/ENW-VICI 865.17.004 (RMHM) and NWO/ENW-XL 2019.029 (MdJ and RMHM), and by Prof. dr. Jan van der Hoevenstichting voor Theoretische Biologie (RMHM) affiliated to the Leiden University Fund (RMHM).

## Supporting information

**Figure S1.**
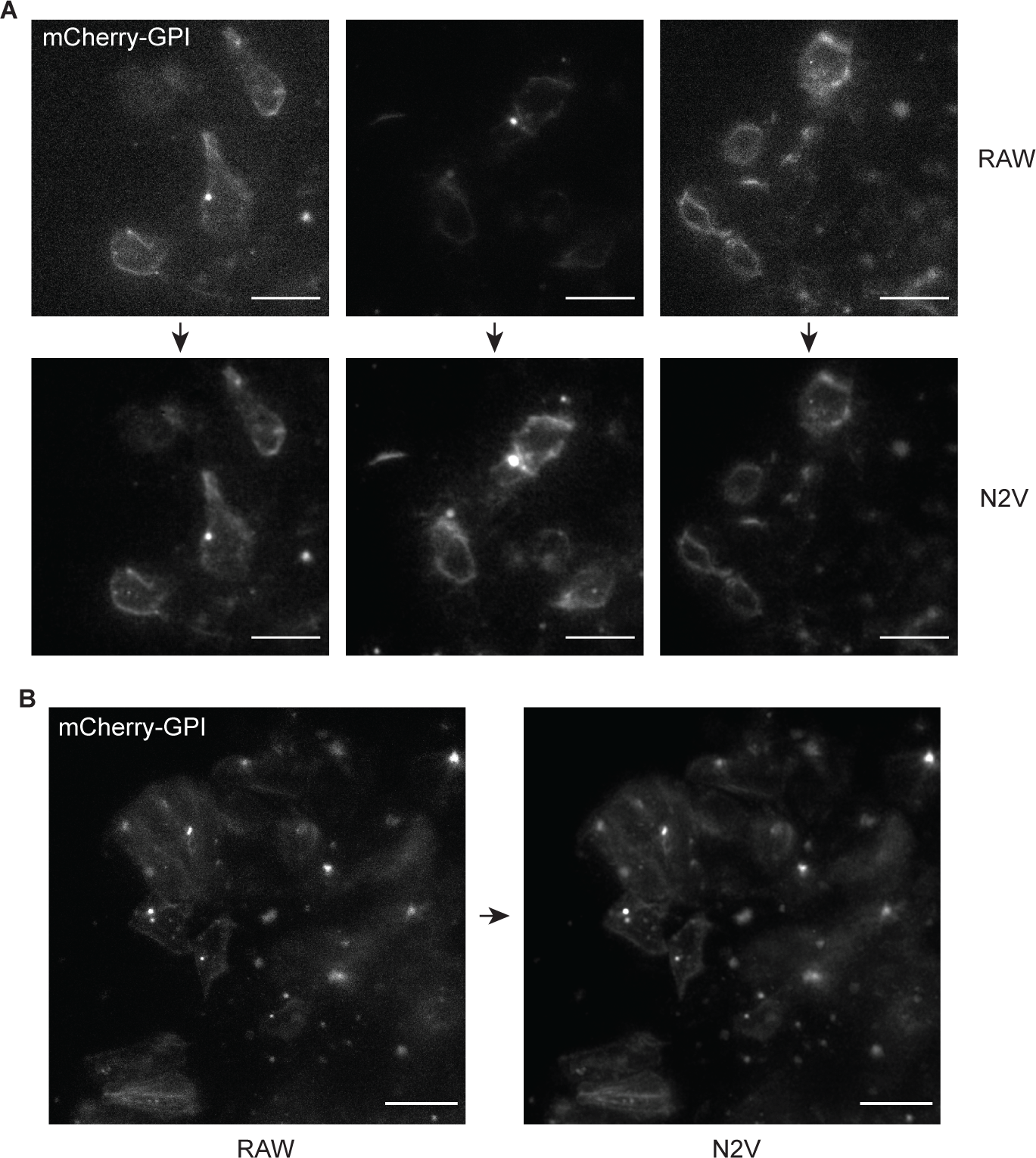
Comparison of raw images and images that were denoised using Noise2Void. A: Image of raw data (top panel) and denoised data (bottom panel) at *t* = 1. A cropped image (505 *×* 505 pixels) is shown of the 11*^th^* plane of the z-stack of the 1*^st^* time point of the time lapse. The denoised image was predicted from the model that was trained on the images of the 1*^st^* time point. B: Image of raw data (top panel) and denoised data (bottom panel) at *t* = 5. A cropped image (505 *×* 505 pixels) is shown of the 10*^th^* plane of the z-stack of the 5*^th^* time point of the time lapse. The denoised image was predicted from the model that was trained on the images of the 7*^th^* time point. C: Image of raw data (top panel) and denoised data (bottom panel) at *t* = 23. A cropped image (505 *×* 505 pixels) is shown of the 6*^th^* plane of the z-stack of the 23*^rd^* time point of the time lapse. The denoised image was predicted from the model that was trained on the images of the 31*^st^* time point, the last time point of the time lapse. D: Image of raw data (left panel) and denoised data (right panel) at *t* = 31. A cropped image (674 *×* 674 pixels) is shown of the maximum projection of the z-stack of the 31*^st^* time point of the time lapse. The denoised image was predicted from the model that was trained on the images of the 31*^st^* time point. A,B,C,D: The minimum displayed value for each image was set to the 5*^th^* percentile of the image intensity distribution to enable a qualitative comparison. Scale bars: 20 *µ*m.

**Figure S2.**
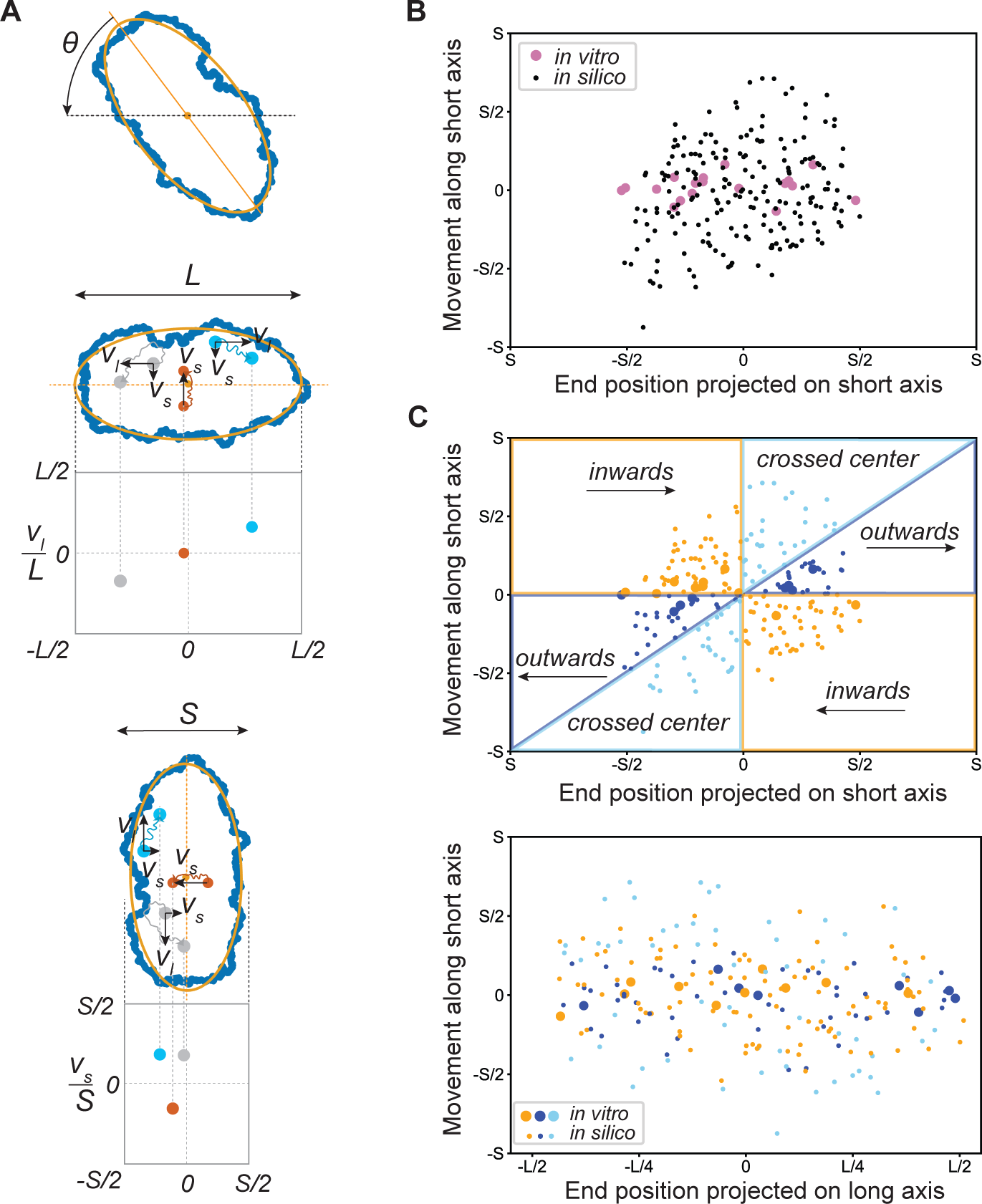
Analysis cell movements with respect to the long and short axis. A: Scheme showing how cell positions were projected on the long axis of the gastruloid. An ellipse was fitted to the outline of the gastruloid. The angle *θ* between the long axis of the ellipse and the nearest orthogonal axis when rotating counter-clockwise was determined for each time point (top panel). Positions of cells were rotated by the angle *θ* to have them projected on the long axis of the gastruloid. Here, the new x coordinate of each cell corresponds to the cell’s position on the long axis, and the new y coordinate to the cell’s position on the short axis. For each cell, the movement along the long axis *v_l_*(*x_end_ − x_start_*) or along the short axis *v_s_* (*y_end_ − y_start_*) was determined, and normalized by the length of the long axis *L* or the short axis *S*, respectively. The resulting *v_l_/L* was plotted against the end position of each cell projected on the long axis (mid panel). The resulting *vs/S* was plotted against the end position of each cell projected on the short axis (bottom panel). B: Distance each cell moved along the short axis plotted against the end position of each cell projected on the short axis. Large, pink circles represent cells of the *in vitro* gastruloid. Small, black circles represent cells of the simulated gastruloid. C: Same plot as depicted in B with cells in the plot being colored by their direction of movement with respect to the long central axis (top panel). Orange: cells are moving inwards, towards the long central axis; dark blue: cells are moving outwards, away from the long central axis; light blue: cells initially move inwards, but eventually move outwards, thereby crossing the long central axis. Bottom panel: distance of each cell moved along the short axis plotted against the end position of each cell projected on the long axis. Cells are colored by their direction of movement with respect to the long central axis. Large circles represent cells of the *in vitro* gastruloid. Small circles represent cells of the simulated gastruloid.

**Figure S3.**
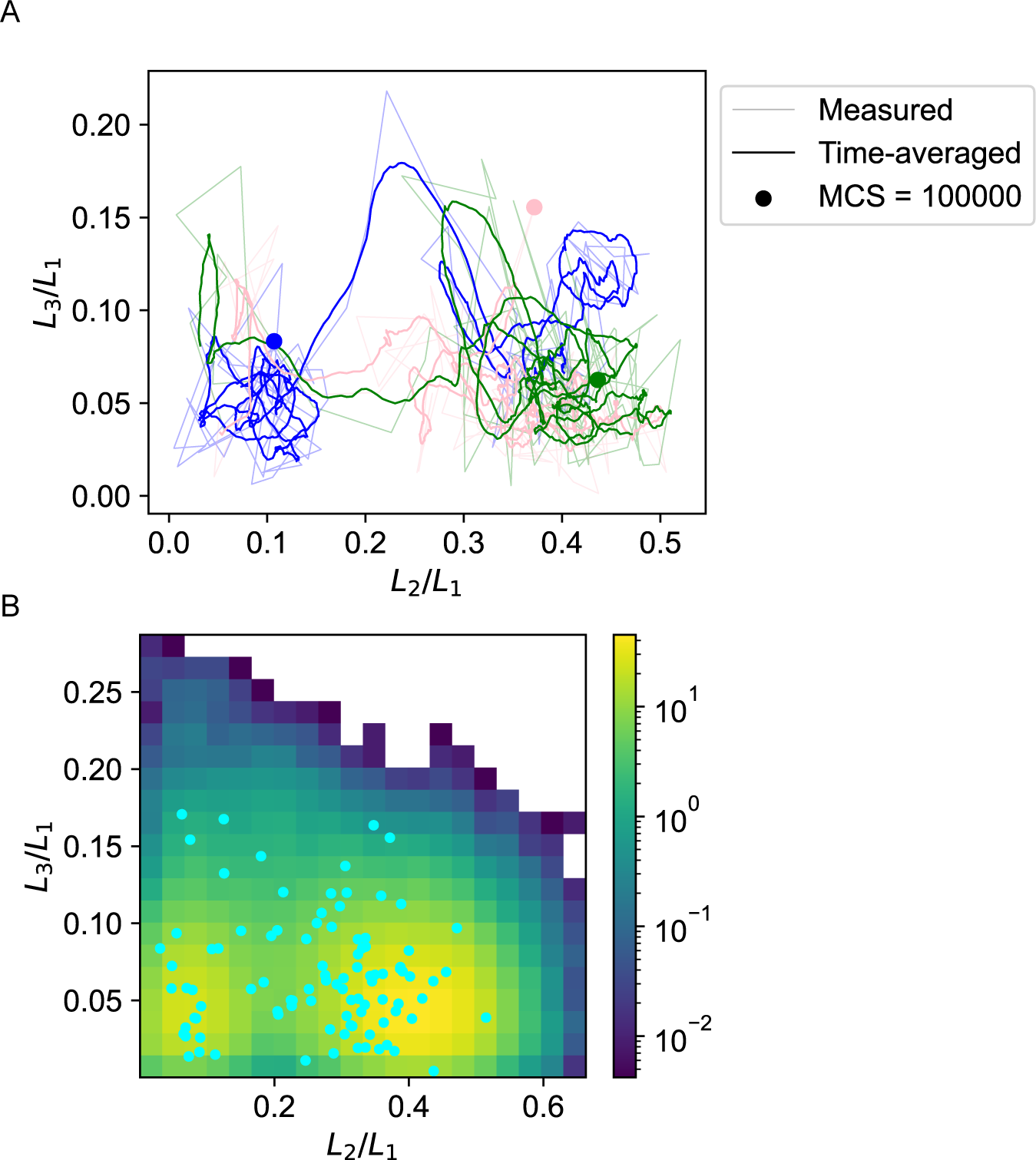
Shape distributions in simulations are not due to fluctuations around local minima. A: The evolution of three simulations with the same parameters (pulling force = 20 and *γ*(*c, M*) = 15) in the *L*_2_*/L*_1_ *− L*_3_*/L*_1_ space is shown over time until 500,000 Monte Carlo steps. The thin lines give the *L*_2_*/L*_1_ and *L*_3_*/L*_1_ values every 5,000 Monte Carlo steps. The thick lines indicate the time averaged value over the last 25 measurements (each 500 Monte Carlo steps apart). The dots highlight the values at 100,000 Monte Carlo steps, which is the simulation time in all other figures. B: Density plot of the *L*_2_ and *L*_3_ values for 100 simulations with the same parameters at different time points, up to 500,000 Monte Carlo steps. The values at 100,000 Monte Carlo steps are indicated with dots.

**Figure S4.**
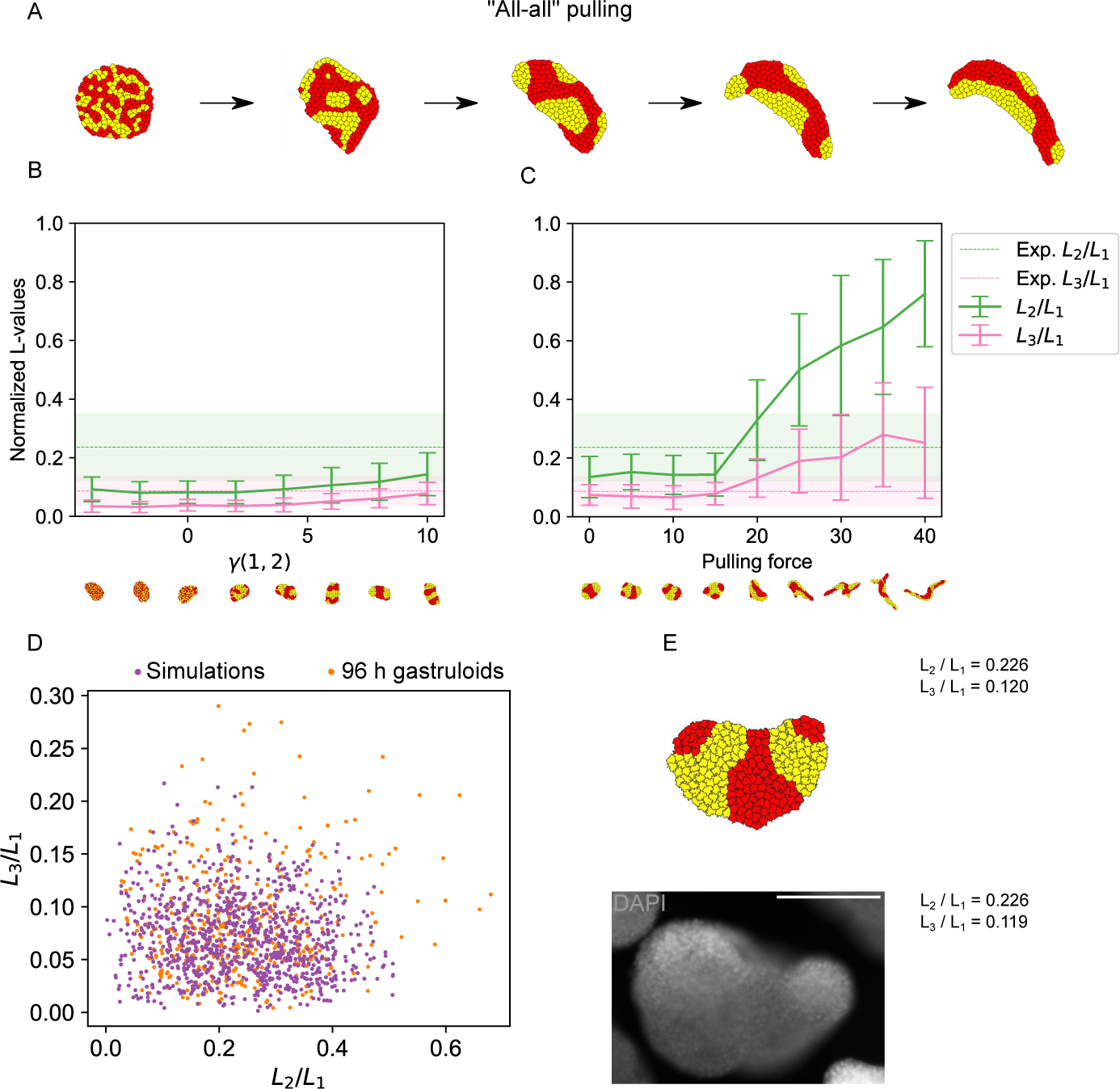
Simulations of CE with two cell types where all cells extend filopodia. A: Example shapes resulting from a simulation of same type pulling for 100, 25,000, 50,000, 75,000 and 100,000 Monte Carlo steps. B,C: Quantification of shapes resulting from simulations of CE with two cell types. Shown are LOCO-EFA coefficients averaged over a population of shapes. The shaded areas (in the experimental data) and vertical bars (in the simulation data) indicate standard deviations. B: The surface tension between the cell types was varied. The pulling force was kept constant at 15. C: The pulling force was varied. The surface tension between the cell types was kept constant at 10. D: Scatter plot of the scaled LOCO-EFA coefficients *L*_3_*/L*_1_ versus *L*_2_*/L*_1_ for the experimental data and simulations of CE with pulling force *λ_F_* = 15 and surface tension *γ*(1, 2) = 10. These parameters gave the smallest 2D Kolmogorov-Smirnov test statistic *D* = 0.37 and *p* = 1.4 . 10*^−^*^7^. E: Example of experimentally observed shape (bottom) with highly similar simulated shape (top) and their corresponding scaled LOCO-EFA coefficients *L*_3_*/L*_1_ and *L*_2_*/L*_1_. Bottom: a 96 h fixed gastruloid. Shown is the mid-plane of a z-stack. Cell nuclei were stained with DAPI. Scale bar: 200 *µ*m.

**Figure S5.**
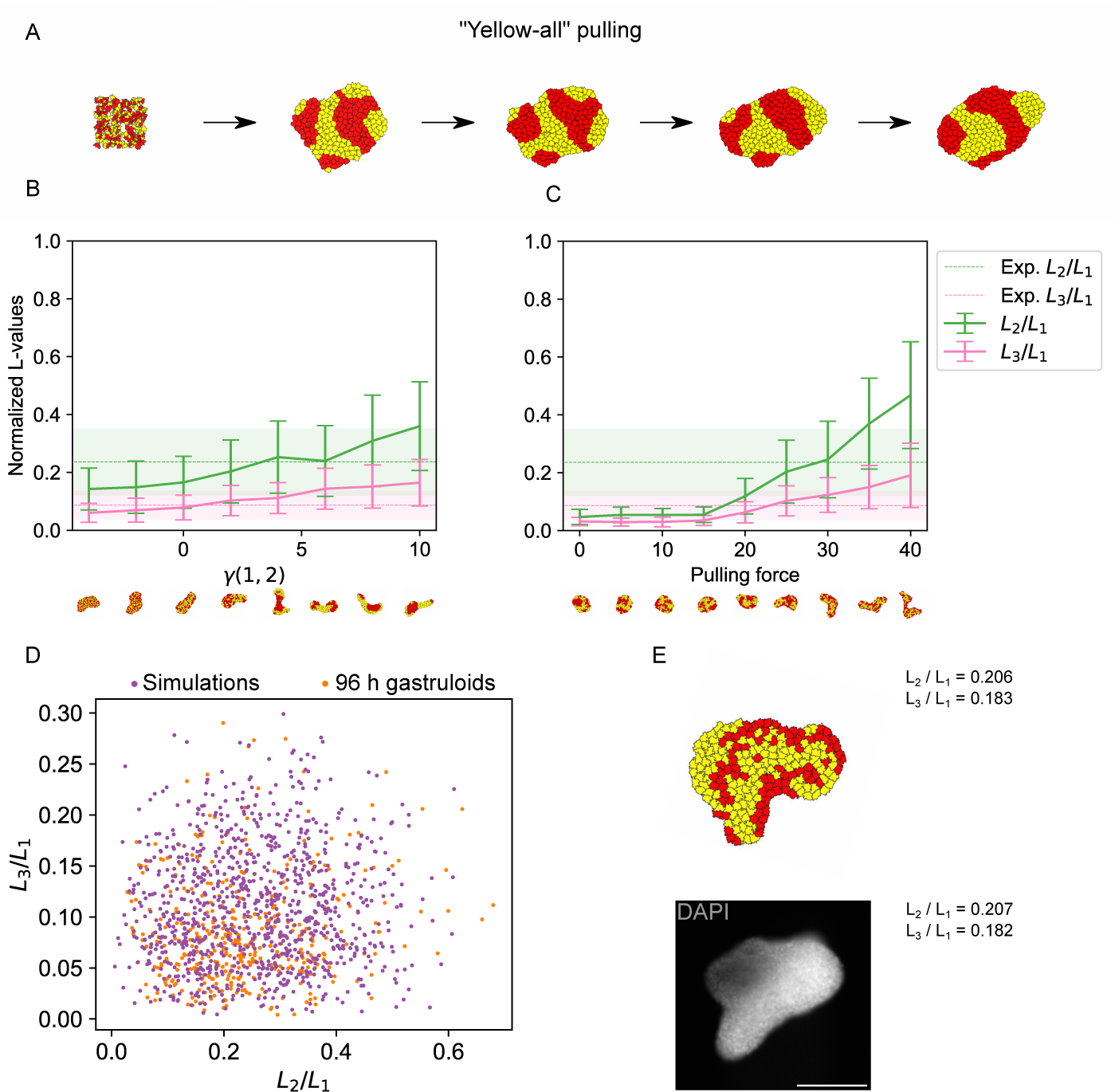
Simulations of CE with two cell types where only yellow cells extend filopodia. A: Example shapes resulting from a simulation of all-all pulling for 100, 25,000, 50,000, 75,000 and 100,000 Monte Carlo steps. B,C: Quantification of shapes resulting from simulations of CE with two cell types. Shown are LOCO-EFA coefficients averaged over a population of shapes. The shaded areas (in the experimental data) and vertical bars (in the simulation data) indicate standard deviations. B: The surface tension between the cell types was varied. The pulling force was kept constant at 25. C: The pulling force was varied. The surface tension between the cell types was kept constant at 2. D: Scatter plot of the scaled LOCO-EFA coefficients *L*_3_*/L*_1_ versus *L*_2_*/L*_1_ for the experimental data and simulations of CE with pulling force *λ_F_* = 25 and interaction energy *γ*(1, 2) = 2. These parameters gave the smallest 2D Kolmogorov-Smirnov test statistic *D* = 0.21 and *p* = 0.0098. E: Example of experimentally observed shape (bottom) with highly similar simulated shape (top) and their corresponding scaled LOCO-EFA coefficients *L*_3_*/L*_1_ and *L*_2_*/L*_1_. Bottom: a 96 h fixed gastruloid. Shown is the mid-plane of a z-stack. Cell nuclei were stained with DAPI. Scalebar: 200 *µm*.

**Figure S6.**
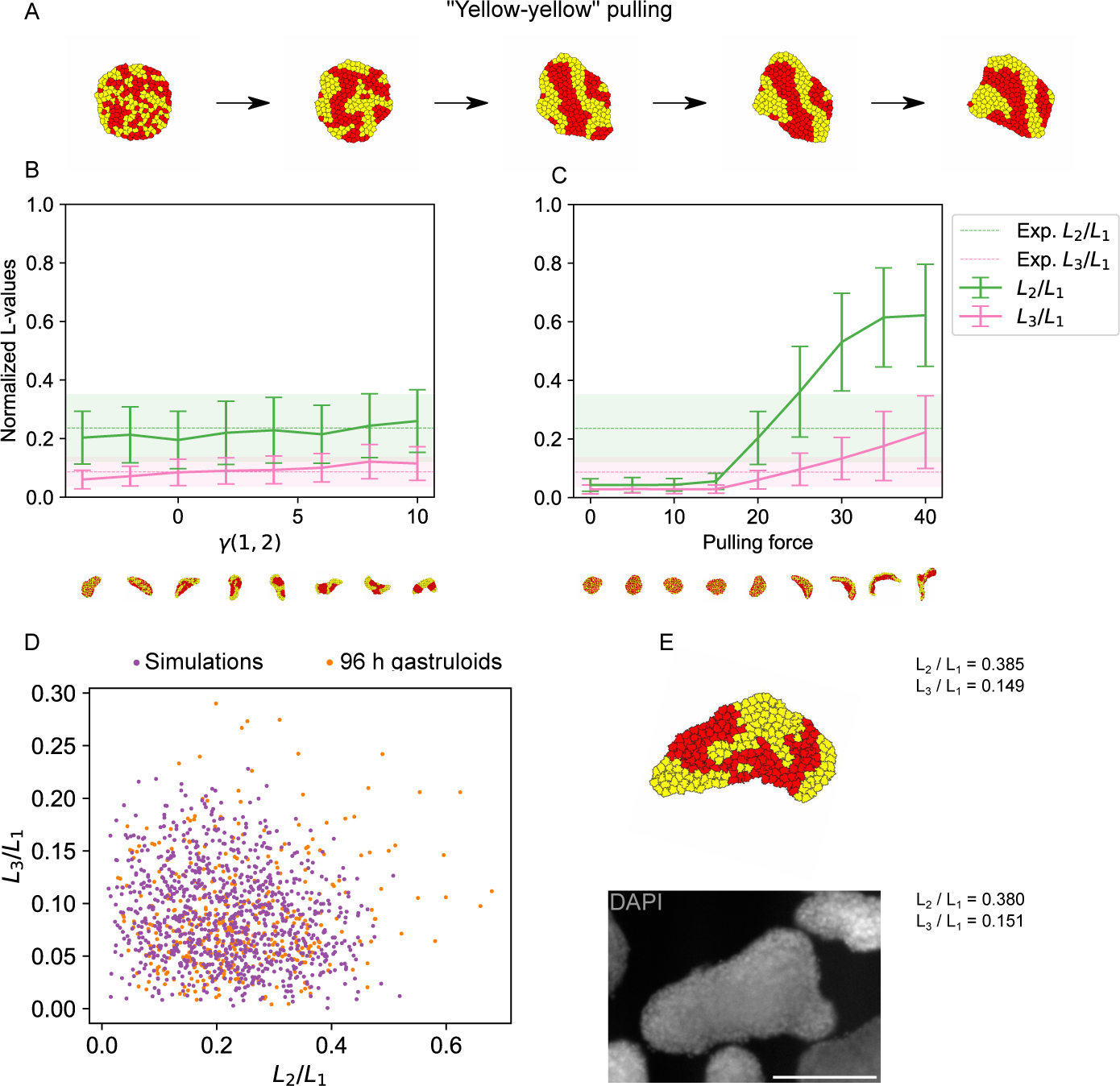
Simulations of CE with two cell types where yellow cells extend filopodia to yellow cells. Red cells do not exert or feel filopodia. A: Example shapes resulting from a simulation of yellow-yellow pulling for 100, 25,000, 50,000, 75,000 and 100,000 Monte Carlo steps. B,C: Quantification of shapes resulting from simulations of CE with two cell types. Shown are LOCO-EFA coefficients averaged over a population of shapes. The shaded areas (in the experimental data) and vertical bars (in the simulation data) indicate standard deviations. B: The surface tension between the cell types was varied. The pulling force was kept constant at 20. C: The surface tension between the cell types was kept constant at 2. D: Scatter plot of the scaled LOCO-EFA coefficients *L*_3_*/L*_1_ versus *L*_2_*/L*_1_ for the experimental data and simulations of CE with pulling force *λ_F_* = 20 and surface tension *γ*(1, 2) = 2. These parameters gave the smallest 2D Kolmogorov-Smirnov test statistic *D* = 0.15 and *p* = 0.12. E: Example of experimentally observed shape (bottom) with highly similar simulated shape (top) and their corresponding scaled LOCO-EFA coefficients *L*_3_*/L*_1_ and *L*_2_*/L*_1_. Bottom: a 96 h fixed gastruloid. Shown is the mid-plane of a z-stack. Cell nuclei were stained with DAPI. Scale bar: 200 *µ*m.

**Figure S7.**
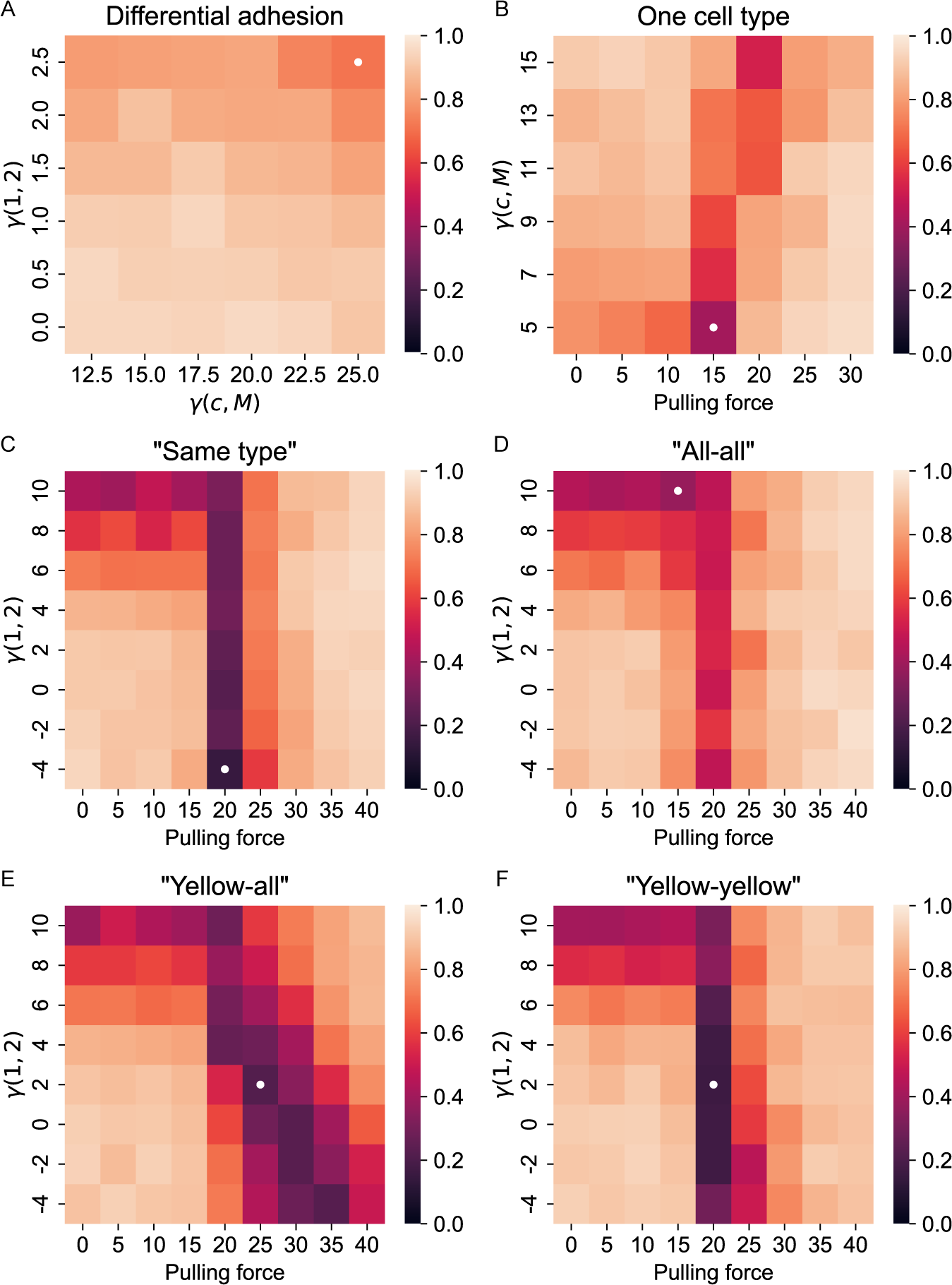
Heat maps of test statistic *D* for different parameters show multiple feasible parameters. The test statistic *D* from the 2D Kolmogorov-Smirnov test has been computed from 100 simulated points for different parameter values and different models. The lowest values are indicated with a white dot. A: For differential adhesion. B: For one cell type pulling. C: For same type pulling. D: For all-all pulling. E: For yellow-all pulling. F: For yellow-yellow pulling.

**Figure S8.**
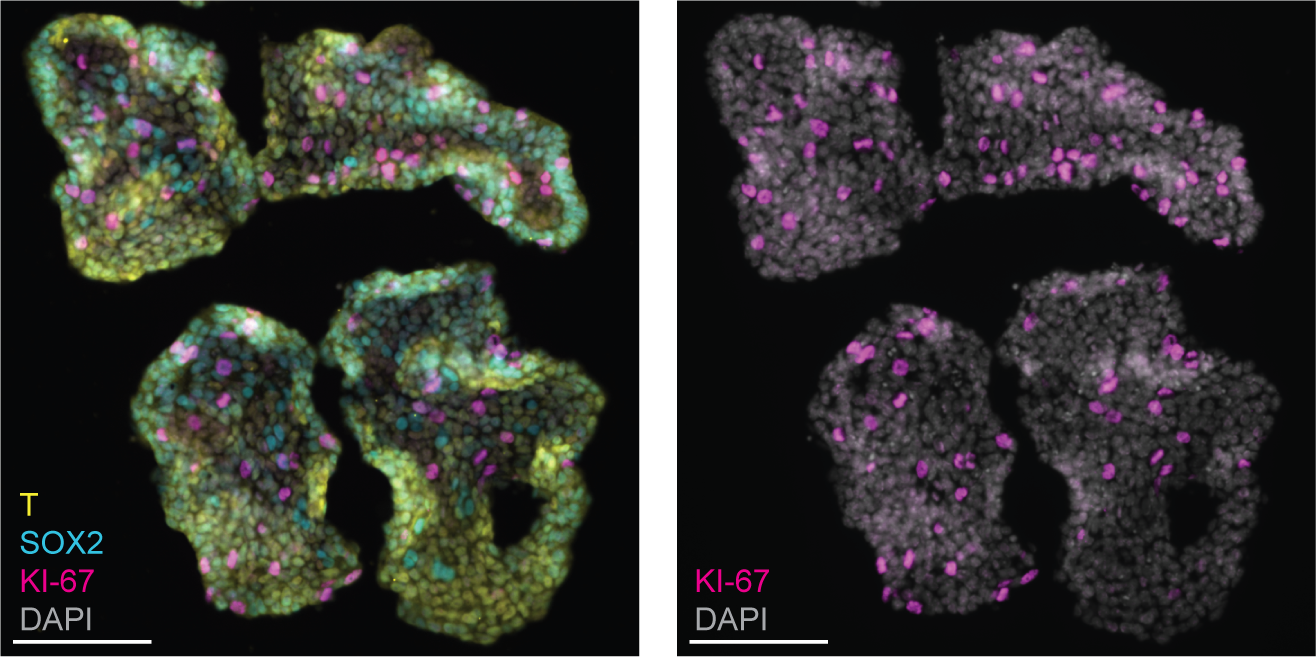
Proliferating cells are found throughout the 96 h gastruloid. Immunostained sections of four gastruloids that were fixed at 96 h. Left: immunostaining of Brachyury/T, Sox2 and the proliferation marker Ki-67. Right: The same sections, but only the immunostaining of Ki-67 is shown. Cell nuclei were stained with DAPI. Scale bars: 100 *µ*m.

**Video 1. Time-lapse lightsheet measurement of elongating gastruloid over 5h. Maximum z-projection. See Fig. 5 for details.**

**Video 2. CPM simulation of elongating gastruloid. See Fig. 5 for details.**

## References

1. McDole K, Guignard L, Amat F, Berger A, Malandain G, Royer LA, et al. In Toto Imaging and Reconstruction of Post-Implantation Mouse Development at the Single-Cell Level. Cell. 2018;175(3):859 – 876.e33. doi:10.1016/j.cell.2018.09.031.

2. Ichikawa T, Zhang HT, Panavaite L, Erzberger A, Fabrèges D, Snajder R, et al. An ex vivo system to study cellular dynamics underlying mouse peri-implantation development. Developmental Cell. 2022;57(3):373–386.e9. doi:10.1016/j.devcel.2021.12.023.

3. Bedzhov I, Zernicka-Goetz M. Self-Organizing Properties of Mouse Pluripotent Cells Initiate Morphogenesis upon Implantation. Cell. 2014;156(5):1032 – 1044. doi:10.1016/j.cell.2014.01.023.

4. van den Brink SC, Baillie-Johnson P, Balayo T, Hadjantonakis AK, Nowotschin S, Turner DA, et al. Symmetry breaking, germ layer specification and axial organisation in aggregates of mouse embryonic stem cells. Development. 2014;141(22):4231 LP – 4242. doi:10.1242/dev.113001.

5. Mongera A, Michaut A, Guillot C, Xiong F, Pourquié O. Mechanics of Anteroposterior Axis Formation in Vertebrates. Annual Review of Cell and Developmental Biology. 2019;35(1):1–25. doi:10.1146/annurev-cellbio-100818-125436.

6. Emig AA, Williams MLK. Gastrulation morphogenesis in synthetic systems. Seminars in Cell & Developmental Biology. 2023;141:3–13. doi:10.1016/j.semcdb.2022.07.002.

7. Zallen JA, Zallen R. Cell-pattern disordering during convergent extension inDrosophila. Journal of Physics: Condensed Matter. 2004;16(44):S5073–S5080. doi:10.1088/0953-8984/16/44/005.

8. Keller R, Danilchik M, Gimlich R, Shih J. The function and mechanism of convergent extension during gastrulation of Xenopus laevis. J Embryol Exp Morphol. 1985;.

9. Heisenberg CP, Tada M, Rauch GJ, Saúde L, Concha ML, Geisler R, et al. Silberblick/Wnt11 mediates convergent extension movements during zebrafish gastrulation. Nature. 2000;405(6782):76–81.

10. Sun Z, Amourda C, Shagirov M, Hara Y, Saunders TE, Toyama Y. Basolateral protrusion and apical contraction cooperatively drive Drosophila germ-band extension. Nature Cell Biology. 2017;19(4):375–383. doi:10.1038/ncb3497.

11. Weng S, Huebner RJ, Wallingford JB. Convergent extension requires adhesion-dependent biomechanical integration of cell crawling and junction contraction. Cell Reports. 2022;39(4):110666. doi:10.1016/j.celrep.2022.110666.

12. Williams M, Yen W, Lu X, Sutherland A. Distinct Apical and Basolateral Mechanisms Drive Planar Cell Polarity-Dependent Convergent Extension of the Mouse Neural Plate. Developmental Cell. 2014;29(1):34–46. doi:https://doi.org/10.1016/j.devcel.2014.02.007.

13. Huebner RJ, Wallingford JB. Coming to Consensus: A Unifying Model Emerges for Convergent Extension. Developmental Cell. 2018;46(4):389–396. doi:10.1016/j.devcel.2018.08.003.

14. Ybot-Gonzalez P, Savery D, Gerrelli D, Signore M, Mitchell CE, Faux CH, et al. Convergent extension, planar-cell-polarity signalling and initiation of mouse neural tube closure. Development. 2007;134(4):789–799. doi:10.1242/dev.000380.

15. Sánchez-Corrales YE, Hartley M, van Rooij J, Marée AFM, Grieneisen VA. Morphometrics of complex cell shapes: lobe contribution elliptic Fourier analysis (LOCO-EFA). Development. 2018;145(6):dev156778. doi:10.1242/dev.156778.

16. Graner F, Glazier JA. Simulation of biological cell sorting using a two-dimensional extended Potts model. Physical Review Letters. 1992;69(13):2013–2016. doi:10.1103/PhysRevLett.69.2013.

17. Glazier J, Graner F. Simulation of the differential adhesion driven rearrangement of biological cells. Physical review E, Statistical physics, plasmas, fluids, and related interdisciplinary topics. 1993;47(3):2128–2154.

18. Zajac M, Jones GL, Glazier JA. Simulating convergent extension by way of anisotropic differential adhesion. Journal of Theoretical Biology. 2003;222(2):247–259. doi:https://doi.org/10.1016/S0022-5193(03)00033-X.

19. Belmonte JM, Swat MH, Glazier JA. Filopodial-Tension Model of Convergent-Extension of Tissues.(Research Article). PLoS Computational Biology. 2016;12(6):e1004952. doi:10.1371/journal.pcbi.1004952.

20. Dekkers JF, Alieva M, Wellens LM, Ariese HC, Jamieson PR, Vonk AM, et al. High-resolution 3D imaging of fixed and cleared organoids. Nature protocols. 2019;14(6):1756–1771.

21. Strnad P, Gunther S, Reichmann J, Krzic U, Balazs B, Medeiros Gd, et al. Inverted light-sheet microscope for imaging mouse pre-implantation development. Nature Methods. 2016;13(2):139–142. doi:10.1038/nmeth.3690.

22. Serra D, Mayr U, Boni A, Lukonin I, Rempfler M, Meylan LC, et al. Self-organization and symmetry breaking in intestinal organoid development. Nature. 2019;569(7754):66–72. doi:10.1038/s41586-019-1146-y.

23. Otsu N. A Threshold Selection Method from Gray-Level Histograms. IEEE Trans Syst Man Cybern. 1979;SMC-9(1):62–66. doi:10.1109/TSMC.1979.4310076.

24. Scharr H. Optimal operators in digital image processing. Heidelberg: Dissertation; 2000.

25. Najman L, Schmitt M. Watershed of a continuous function. Signal Processing, Elsevier. 1994;38(1):99–112. doi:10.1016/0165-1684(94)90059-0.

26. Schindelin J, Arganda-Carreras I, Frise E, Kaynig V, Longair M, Pietzsch T, et al. Fiji: an open-source platform for biological-image analysis. Nature Methods. 2012;9(7):676–682. doi:10.1038/nmeth.2019.

27. Krull A, Buchholz TO, Jug F. Noise2Void - Learning Denoising From Single Noisy Images. In: Proceedings of the IEEE/CVF Conference on Computer Vision and Pattern Recognition (CVPR); 2019.

28. Bradski G. The OpenCV Library. Dr Dobb’s Journal of Software Tools. 2000;.

29. Walt Svd, Schönberger JL, Nunez-Iglesias J, Boulogne F, Warner JD, Yager N, et al. scikit-image: image processing in Python. PeerJ. 2014;2:e453. doi:10.7717/peerj.453.

30. Merks RMH, Glazier JA. A cell-centered approach to developmental biology. Physica A: Statistical Mechanics and its Applications. 2005;352(1):113–130. doi:https://doi.org/10.1016/j.physa.2004.12.028.

31. Daub JT, Merks RM. Cell-based computational modeling of vascular morphogenesis using Tissue Simulation Toolkit. In: Vascular morphogenesis. Springer; 2015. p. 67–127.

32. Tange O. Gnu parallel-the command-line power tool. USENIX Magazine. 2011;36(1):42–47.

33. Fasano G, Franceschini A. A multidimensional version of the Kolmogorov–Smirnov test. Monthly Notices of the Royal Astronomical Society. 1987;225(1):155–170.

34. Ness-Cohn E, Braun R. Fasano-Franceschini Test: an Implementation of a 2-Dimensional Kolmogorov-Smirnov test in R. arXiv preprint arXiv:210610539. 2021;.

35. Bérenger-Currias NMLP, Mircea M, Adegeest E, van den Berg PR, Feliksik M, Hochane M, et al. Extraembryonic endoderm cells induce neuroepithelial tissue in gastruloids. bioRxiv. 2021;doi:10.1101/2020.02.13.947655.

36. Vroomans RMA, Hogeweg P, ten Tusscher KHWJ. Segment-Specific Adhesion as a Driver of Convergent Extension. PLOS Computational Biology. 2015;11(2):e1004092.

37. Yen WW, Williams M, Periasamy A, Conaway M, Burdsal C, Keller R, et al. PTK7 is essential for polarized cell motility and convergent extension during mouse gastrulation. Development. 2009;136(12):2039–2048. doi:10.1242/dev.030601.

38. Marikawa Y, Tamashiro DAA, Fujita TC, Alarcón VB. Aggregated P19 mouse embryonal carcinoma cells as a simple in vitro model to study the molecular regulations of mesoderm formation and axial elongation morphogenesis. genesis. 2009;47(2):93–106. doi:10.1002/dvg.20473.

39. Shawlot W, Deng JM, Behringer RR. Expression of the mouse cerberus-related gene, Cerr1, suggests a role in anterior neural induction and somitogenesis. Proceedings of the National Academy of Sciences. 1998;95(11):6198–6203. doi:10.1073/pnas.95.11.6198.

40. Beccari L, Moris N, Girgin M, Turner DA, Baillie-Johnson P, Cossy AC, et al. Multi-axial self-organization properties of mouse embryonic stem cells into gastruloids. Nature Publishing Group. 2018;5:1. doi:10.1038/s41586-018-0578-0.

41. Hashmi A, Tlili S, Perrin P, Lowndes M, Peradziryi H, Brickman JM, et al. Cell-state transitions and collective cell movement generate an endoderm-like region in gastruloids. eLife. 2022;11:e59371. doi:10.7554/elife.59371.

42. Semrau S, Idema T, Holtzer L, Schmidt T, Storm C. Accurate determination of elastic parameters for multicomponent membranes. Physical Review Letters. 2008;100(8):088101.

43. Cermola F, D’Aniello C, Tatè R, Cesare DD, Martinez-Arias A, Minchiotti G, et al. Gastruloid Development Competence Discriminates Different States of Pluripotency. Stem Cell Reports. 2021;16(2):354–369. doi:10.1016/j.stemcr.2020.12.013.

44. Mantziou V, Baillie-Benson P, Jaklin M, Kustermann S, Arias AM, Moris N. In vitro teratogenicity testing using a 3D, embryo-like gastruloid system. Reproductive Toxicology (Elmsford, Ny). 2021;105:72–90. doi:10.1016/j.reprotox.2021.08.003.

