## Supplementary figures and images for "A combination of convergent extension and differential adhesion explains the shapes of elongating gastruloids"

### Video2

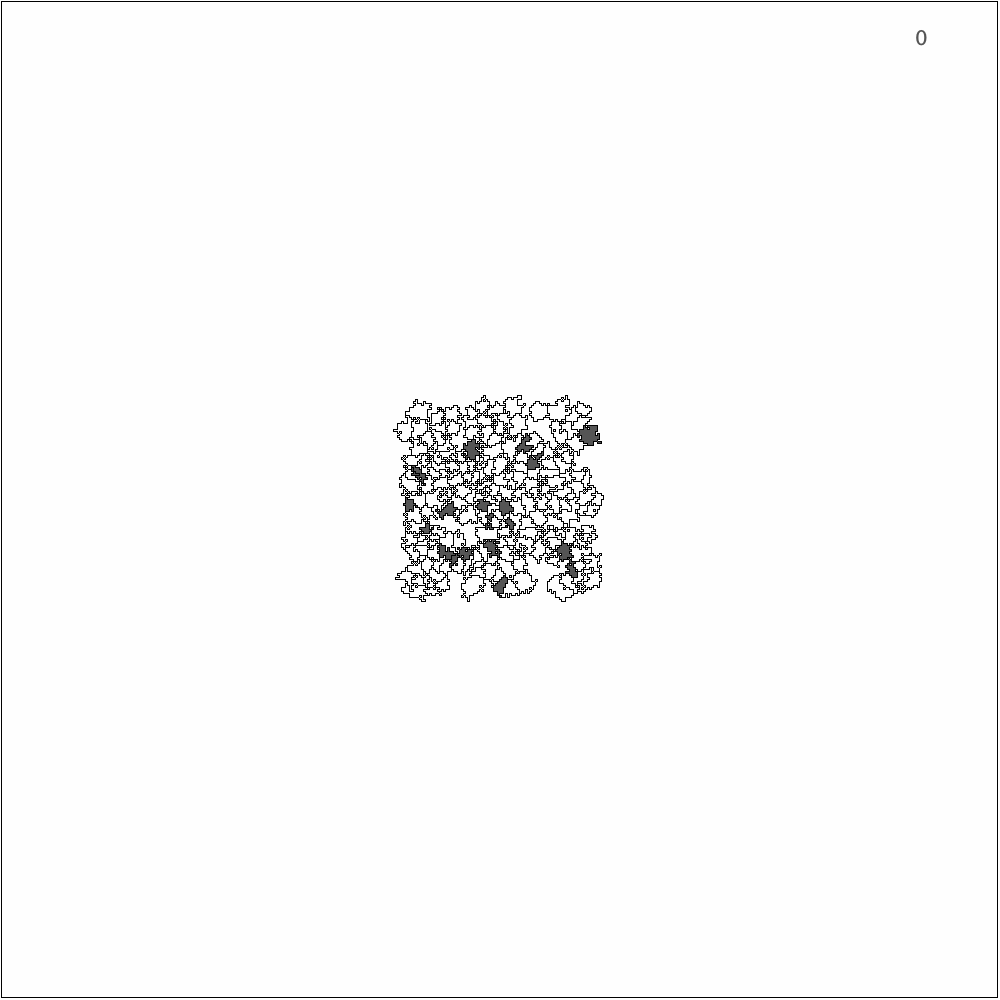
